# A Druggable G Protein Checkpoint in Cholesterol Efflux

**DOI:** 10.64898/2026.02.15.705962

**Authors:** Gajanan D. Katkar, Mahitha Shree Anandachar, Celia R. Espinoza, Megan Estanol, Caroline Biggs, Michelle Nacayama, Shu-Tsing-Hsu, Ellie Tam, Raktim Mukherjee, Ella McLaren, Stephanie Aviles, Vanessa Castillo, Yashaswat S. Malhotra, Madhubanti Mullick, Armando Aaron, Oswald Quehenberger, Jerry Yang, Benedikt Kaufmann, Saptarshi Sinha, Pradipta Ghosh

## Abstract

Immunometabolic diseases such as obesity, fatty liver, and atherosclerosis arise when lipid-associated macrophages (LAMs) fail to resolve lipid overload via reverse cholesterol transport (RCT), the body’s sole pathway for lipid disposal. How RCT is restrained in disease remains unknown. Integrating systems modeling with human plaque transcriptomes, we identify LAM subpopulations associated with plaque progression and uncover *CCDC88A* (GIV) as a macrophage-intrinsic checkpoint on RCT. Myeloid-specific GIV deletion in mice reduces aortic plaque burden, mobilizes hepatic and adipose lipid stores and promotes fecal sterol disposal. Mechanistically, GIV sequesters the cholesterol transporter ABCA1 within endomembranes and activates Gαi●βγ signaling to suppress cAMP/PKA–CREB-dependent efflux programs. Pharmacogenomic disruption of this checkpoint reactivates efflux programs. In human plaque-in-a-dish models targeting this pathway restored efflux where statins and β-blockers failed, translating to an estimated ∼98% reduction in plaque-progression risk in outcome modeling. Thus, RCT-restoration represents a macrophage-intrinsic therapeutic paradigm for immunometabolic disease.

**GRAPHIC ABSTRACT:** 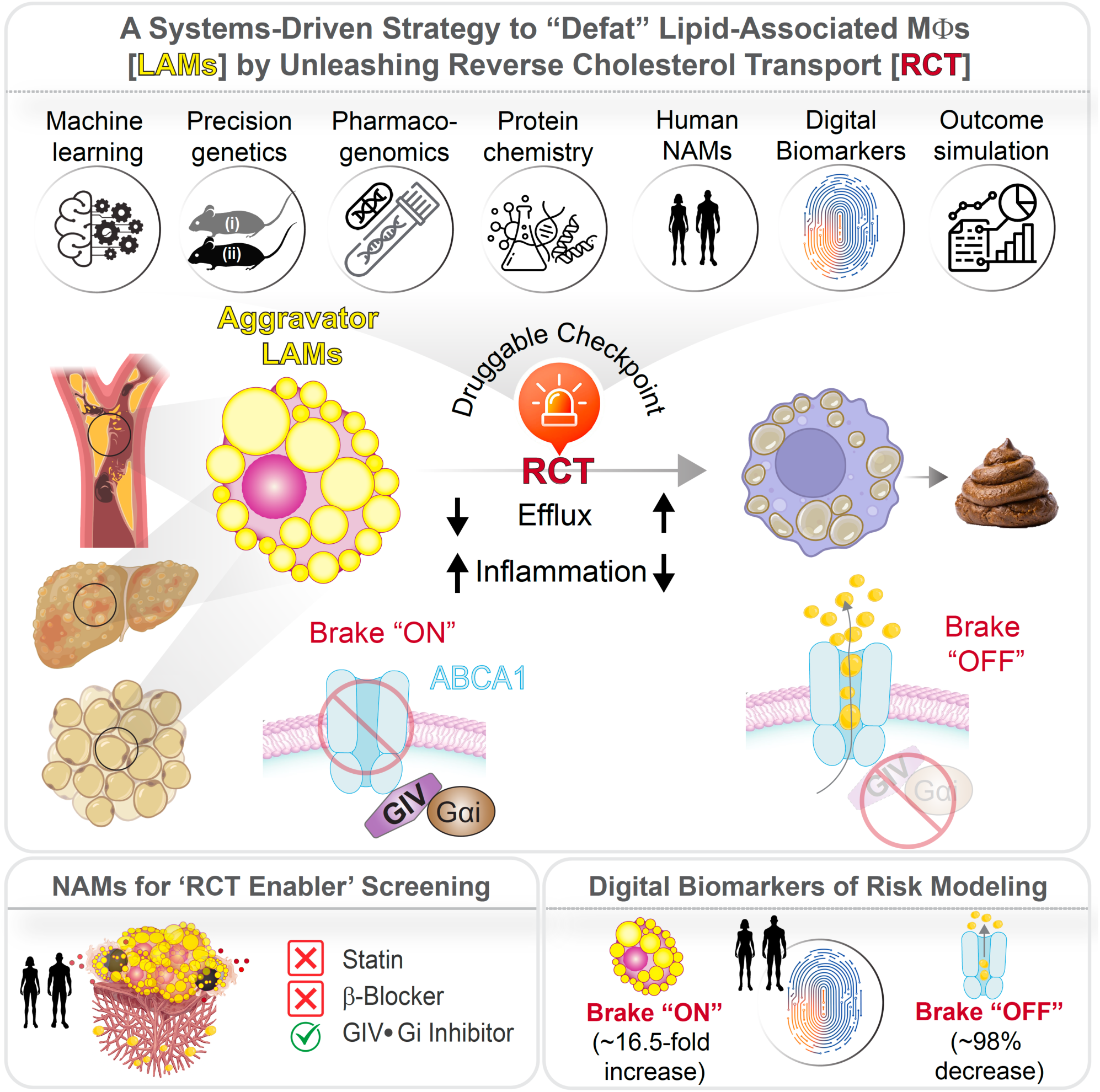

**eTOC blurb:** Lipid-associated macrophages drive immunometabolic disease. *Katkar et al.* show that disabling a GIV-dependent G-protein brake restores cholesterol efflux, reverses plaque lipid accumulation, and establishes reverse cholesterol transport as a druggable therapeutic axis.

**Highlights:**

- Statins slow but rarely reverse plaque burden, leaving residual risk driven by LAM dysfunction
- GIV (CCDC88A) non-canonically modulates Gαi to suppress macrophage cholesterol efflux
- GIV loss or inhibition restores ABCA1 activity via transcriptional and post-translational control
- Blocking GIV●Gαi checkpoint defats LAMs, regresses plaques, and relieves systemic lipid overload
- SIdentifies a druggable node that redefines RCT restoration as a therapeutic paradigm in immunometabolic disease

## INTRODUCTION

Metabolic diseases such as obesity, fatty liver, and atherosclerosis share a single pathological bottleneck that drives disease progression: tissue-resident macrophages that become lipid-engorged, transform into ‘foamy’ cells, and retain excess harmful cholesterol within tissues (reviewed in^1–5^). Approved and emerging therapies, e.g., dietary interventions, statins, or lipoprotein(a)-directed agents, primarily reduce circulating lipids, but cannot evacuate lipids already sequestered within macrophages and other tissue-resident immune cells, leaving substantial residual cardiometabolic risk.

The body’s sole pathway for clearing these intracellular lipid stores is macrophage-to-feces reverse cholesterol transport (**RCT**)^6^. RCT begins with efflux through ATP-binding cassette transporters ABCA1 (ATP-binding cassette transporter A1) and ABCG1 (ATP-binding cassette transporter G1) to high-density lipoprotein (HDL), which delivers lipids to the liver for biliary excretion. In fact, HDL’s cardioprotective effect is attributed in part to its capacity to mobilize excess cholesterol from artery-wall macrophages^7,8^. Among these transporters, ABCA1 serves as the central coordinator of and rate-limiting node of RCT, governed by cAMP–protein kinase A (PKA) signaling^9–14^ and its transcriptional effector CREB^15^. Although cholesterol efflux and elevated cAMP/CREB activity are known to be strongly atheroprotective^15–20^, the upstream molecular regulators that restrain this axis remain poorly defined^21^, representing a major barrier to therapeutically reactivating RCT.

Single-cell RNA sequencing has revealed striking macrophage heterogeneity within lipid-rich tissues^3,22,23^, leading to the identification of lipid-associated macrophages (**LAMs**)^1,3,24–27^, a specialized subset of macrophages tasked with handling lipid overload. First described in adipose depots during obesity^24^, through a defining surface marker, TREM2⁺ (triggering receptor expressed on myeloid cells 2), LAMs were initially viewed as relatively passive lipid buffers that sequester excess cholesterol to preserve tissue homeostasis^24^. Genetic ablation of Trem2 established TREM2 signaling as a central axis in metabolic immunity^24^; however, small-molecule agonists subsequently revealed that its role in LAMs is more complex than initially interpreted^28,29^. In advanced lesions, TREM2 enhances macrophage survival, cholesterol efflux, and efferocytosis, curbing necrosis and inflammation, and stabilizing and halting lesion growth^29,30^, whereas in early lesions, TREM2-driven lipid uptake accelerates lesion growth^28,29^. This biphasic behavior reflects a broader disease trajectory--from early tolerogenic programs that stabilize lipid-laden macrophages and promote plaque expansion, to later inflammatory transitions that drive necrotic core formation^31^. It also underscores that LAMs span a functional spectrum, capable of both *aggravating* and *alleviating* disease by exhibiting a functional spectrum that can vary depending on disease state. These dynamics caution against indiscriminate targeting of LAMs and instead point to upstream regulation of their lipid-handling capacity, specially RCT, as a more rational therapeutic entry point.

Despite its centrality, clinical efforts to harness RCT have repeatedly failed^32–34^. Decades of ApoA1-raising strategies, including ApoA-1 Milano^35,36^, CER-001^37^, and the AEGIS-II trial^38–40^, showed little efficacy in reversing plaque burden. These disappointments highlight a critical knowledge gap: while RCT is the sole mechanism for lipid clearance, the molecular “brakes” that constrain it within tissues remain unknown (**Figure 1A**). Identifying and disabling such brakes could enable not only regression of atherosclerotic plaques but also defatting of the liver and restoration of systemic lipid balance.

**Figure 1:**
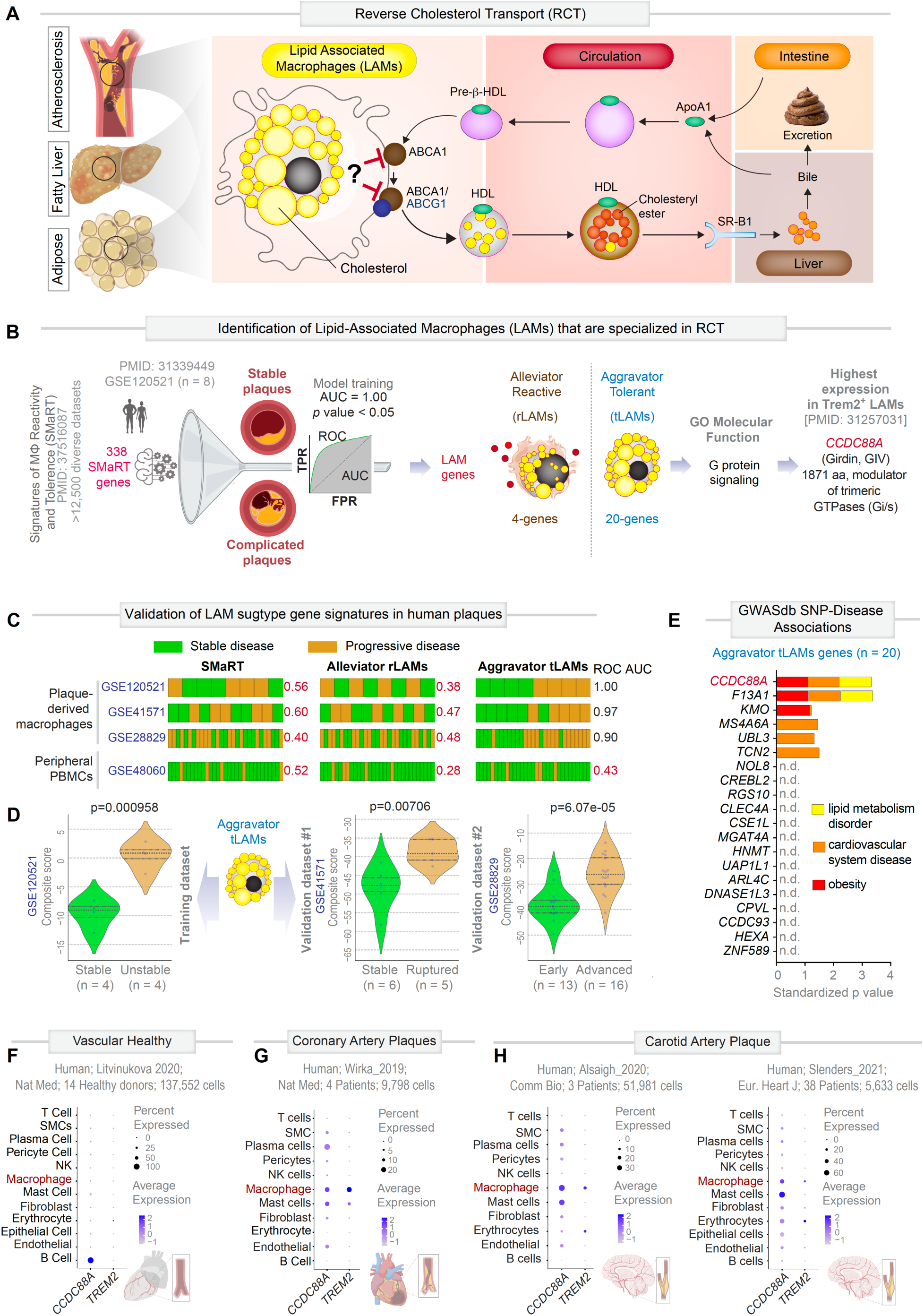
Identification of *CCDC88A* as a checkpoint regulator of lipid-associated macrophages. **A.** Schematic of the RCT pathway, highlighting the role of LAMs in mediating cholesterol efflux from peripheral tissues (vascular, liver, and adipose) via ABCA1/ABCG1 transporters. **B.** *<u>Left</u>*: Strategy to identify subpopulations of aggressor and alleviator LAMs using the previously published computational model of macrophage continuum states, *<u>S</u>*ignatures of *<u>Ma</u>*crophage *<u>R</u>*eactivity and *<u>T</u>*olerance (SMaRT^44^) [see gene list in **Data S1 and Figure S1A-S1D**]. SMaRT was trained on RNAseq datasets from stable versus complicated carotid plaques (GSE120521; n = 8). AUC analysis (AUC = 1.0, *p* < 0.05) distilled LAM-associated SMaRT genes into a separate set of alleviator reactive LAM genes (rLAMs, n = 4) and aggravator tolerant LAM genes (tLAMs, n = 20) [see gene list in **Table S1**]. Gene Ontology analysis revealed enrichment of aggravator LAMs in G protein signaling. Notably, only one gene—*CCDC88A* (Girdin)—was shared with *TREM2⁺* LAMs^24^ [see **Data S2**]. **C.** Validation of rLAM and tLAM gene signatures across independent datasets from human plaques (GSE120521, GSE41571, GSE28829) and PBMCs (GSE48060). Bar plots show the distribution of stable (green) vs. progressive (beige) disease samples within each cohort, rank ordered based on the composite gene score using either the SMaRT (left) or the SMaRT-derived LAM signatures (rLAM, middle; and tLAM, right) signatures from panel B. ROC AUC values indicate the performance of signature classification across datasets. **D.** Violin plots of composite tLAM gene scores in stable versus progressive plaques across training (GSE120521) and independent validation datasets (GSE41571 and GSE28829). Significance assessed by Welch’s t-test. **E.** GWAS enrichment analysis of tLAM genes (n = 20), highlighting strong associations with lipid metabolism disorders (yellow), cardiovascular diseases (orange), and obesity (red). Bars represent standardized *p*-values. **F–H**. Single-cell transcriptomic analysis of *CCDC88A* and *TREM2* across vascular macrophages (from plaqview: https://plaqviewv2.pods.uvarc.io/): (F) healthy vasculature (Litvinukova et al., 2020)^46^, (G) coronary artery plaques (Wirka et al., 2019)^47^, and (H) carotid artery plaques (Alsaigh et al., 2020^48^ and Slenders et al., 2021^49^). Dot plots show fraction of cells (% cells; dot size) expressing each gene and average normalized expression (color intensity).

Here, we identify one such brake. Using systems-level approaches, we formally define the functional spectrum of LAMs—both aggravators and alleviators—and uncover GIV (Gα-interacting vesicle-associated protein, a.k.a Girdin; encoded by *CCDC88A*) as a key facilitator of LAM genesis. Functional and pharmacogenomic studies reveal that GIV-dependent trimeric GTPase activation suppresses cAMP/PKA-CREB signals^41–43^, enforcing an intracellular checkpoint that restrains ABCA1-dependent efflux in aggravator LAMs. Disrupting this axis releases the brake, reactivates RCT and drives macrophage defatting across vascular, hepatic and adipose tissues during systemic cholesterol overload. Together, this work reframes macrophages from passive foam cells into programmable effectors of lipid clearance, establishing restoration of RCT in LAMs as a new therapeutic paradigm for immunometabolic disease.

## RESULTS

### Systems-level dissection of LAMs identifies *CCDC88A* as a putative lipid-handling checkpoint

To resolve functional heterogeneity within lipid-associated macrophages (LAMs, **Figure S1A**), we adopted a two-step systems-level strategy that integrates cohort-scale transcriptomics with orthogonal genetic and single-cell validation.

First, we leveraged our previously developed computational framework, *<u>S</u>*ignatures of *<u>Ma</u>*crophage *<u>R</u>*eactivity and *<u>T</u>*olerance (SMaRT)^44^, which models macrophage states along inflammatory and tolerogenic continua (**Figure 1B, Figure S1B-D, Data S1**). Using bulk RNA-seq datasets from stable and complicated human carotid plaques (GSE120521), we retrained SMaRT to identify transcripts that best distinguish disease states. This unbiased approach distilled a minimal set of 24 LAM-associated gene signature that achieved perfect classification performance (AUC = 1.0, p < 0.05). These genes segregated into two functionally distinct modules: alleviator reactive LAMs (**rLAMs**; n = 4) that are enriched in stable plaques, and aggravator tolerant LAMs (**tLAMs**; n = 20) that are enriched in unstable/complicated plaques (**Figure 1B, Table S1**).

Next, we prioritized candidate regulators within these modules through ontology-guided filtering and cohort-level anchoring. Gene Ontology analysis identified G-protein signaling as the dominant pathway within the disease-associated tLAM module, suggesting at signaling checkpoints that could constrain lipid-handling and or inflammation. Among these candidates, *CCDC88A,* encoding the noncanonical G-protein activator GIV (a.k.a. Girdin) stood out as the only highly expressed transcript shared with canonical TREM2⁺ LAMs (**Figure 1B, Figure S1E, Data S2**). Notably, independent genome-wide functional-genomic screening^45^ identified *CCDC88A* depletion as a modifier of macrophage state transitions linked to lipid metabolism (**Figure S1F**), further prioritizing GIV as a candidate checkpoint in progressive atherosclerosis.

We next validated the discriminatory capacity of these alleviator and aggravator LAM signatures across 3 independent human cohorts, comprised of 44 samples. Aggravator tLAM (aggravator), but not alleviator rLAM signature, robustly stratified samples into stable versus complicated disease states with high fidelity (**Figure 1C**). Composite aggravator LAM scores were significantly higher in complicated plaques compared to their stable counterparts across all datasets (p < 0.05; **Figure 1D**), reinforcing their association with disease progression.

Genome-wide association study (GWAS) anchored these findings at the population level, linking the aggravator tLAM module to lipid disorders, obesity, and cardiovascular disease (**Figure 1E, Figure S1G-H**). Finally, single-cell transcriptomic mapping across human vascular tissues (59 single-cell RNA-seq samples across four independent datasets) revealed that *CCDC88A* is selectively expressed, and often co-expressed with TREM2, in plaque macrophages^46^—including both coronary^47^and carotid^48,49^ lesions, but is largely absent from macrophages in healthy vessels (**Figure 1F-H**).

Collectively, these orthogonal analyses—from systems modeling to cohort validation, human genetics, and single-cell mapping—identify *CCDC88A* (GIV) as a defining molecular node of aggravator disease-modifying LAMs, nominating it as a candidate signaling checkpoint that may constrain macrophage lipid handling and inflammation.

### Myeloid GIV (*CCDC88A*) restricts RCT and promotes lipid accumulation in tissues

GIV is a multimodular signal transducer and the prototypical member of the non-receptor *<u>G</u>*uanine nucleotide *<u>E</u>*xchange *<u>M</u>*odulator (GEM^41^) family of proteins. Unlike the canonical GPCR/G protein pathway, in which G proteins engage exclusively with ligand-activated GPCRs, GEMs like GIV bind and modulate G protein activity downstream of a myriad of cell-surface receptors^20,42,50^. Of relevance here, GIV is a ubiquitously expressed molecule that is highly expressed in immune cells such as macrophages and serves as a signal transducer downstream of PRRs, TLR4^50^, and NOD2, and its gene (*CCDC88A*) is a key determinant of macrophage polarization in the SMaRT model^44^.

To test whether myeloid GIV regulates systemic lipid flux, we subjected myeloid-specific GIV-KO mice^50^ to diet-induced hyperlipidemia (see *Methods*; **Figure 2A**). GIV loss reduced atherosclerotic plaque burden by ∼57%, as revealed by *en face* Oil Red O (ORO) staining of aortas (**Figure 2B-C**), a phenotype reproduced in hyperlipidemic ApoE^⁻/⁻^ backgrounds (AKO vs. DKO; see *Methods*) (**Figure S2A-C**). Beyond the vasculature, hepatic steatosis was attenuated: liver sections from myeloid-specific GIV-deficient mice displayed fewer and smaller lipid vacuoles (“fat balloons”; **Figure 2D-E**). In adipose tissue, loss of myeloid-specific GIV mitigated adipocyte hypertrophy (**Figure 2F and 2H**) and reduced F4/80⁺ macrophage infiltration and crown-like structures^51^, hallmarks of adipose inflammation (**Figure 2G, 2I** and **2J**). Notably, these tissue-level benefits occurred without changes in circulating lipid levels, blood myeloid cell abundance (**Figure S2D-J**), body weight, or organ mass (**Figure S3**), suggesting that GIV’s role may be due to myeloid-intrinsic lipid handling rather than systemic lipid exposure.

**Figure 2.**
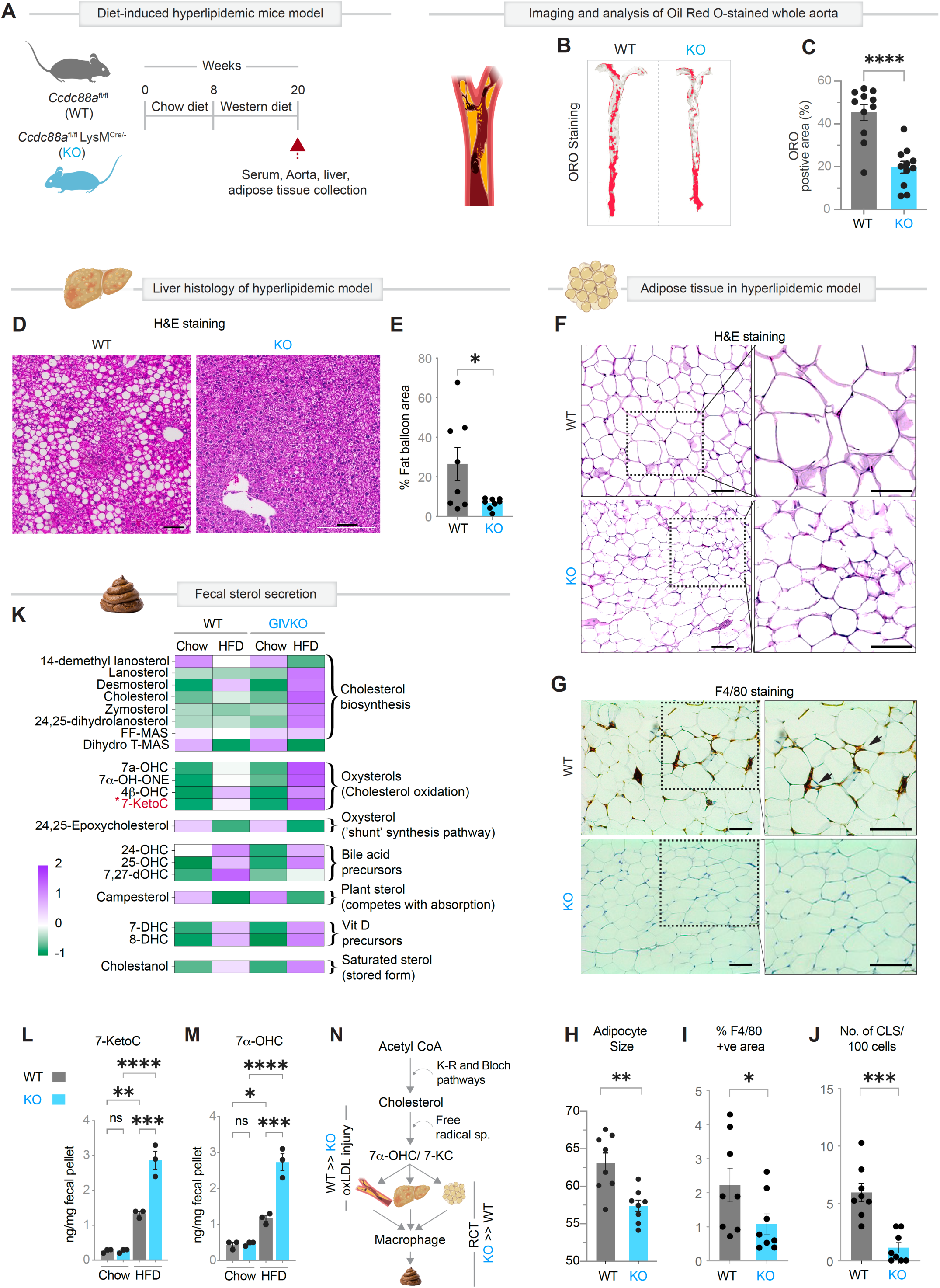
Myeloid GIV loss alleviates plaque burden and mobilizes lipids via enhanced RCT. **A.** Experimental design: Eight-week-old mice were fed a high-fat diet (HFD) for 12 weeks prior to collection of tissues (aorta, serum, and heart; n=11). Data represents two independent cohorts. See also **Figure S2 and S3** for metabolic, hematologic and anatomical parameters and **Figure S2A-C** orthogonal readouts in a genetic (*ApoE*^-/-^) and diet-induced model. **B-C.** Plaque burden: Representative *en face* aortic preparations stained with Oil Red O (B) and quantification of lesion area (C, n=11). Scale bar, 2.5 mm. **D-E.** Liver lipid accumulation: H&E-stained liver sections (D) and quantification of macrovesicular “fat ballooning” (E, n=8). Scale bar, 50 µm. **F-J.** Adipose tissue inflammation: H&E-stained adipose tissue (F) with adipocyte size quantification (H, n=8), Scale bar, 50 µm. F4/80 immunostaining of adipose tissue (representative fields in G) with quantification of F4/80+ area (I) and crown-like structures per 100 adipocytes (J, n=8). Scale bar, 50 µm. **K-M.** Enhanced reverse cholesterol transport: Heatmap of fecal sterol profiles (K) from WT and KO mice either fed chow or high-fat diet (HFD). Fecal pellets were collected between weeks 4-8 of diet exposure. Quantification of 7-KetoC (L) and 7-αOHC (M). See also, **Figure S4** for quantification of various targeted fecal sterols (n=3). **N**. Model: Working model summarizing how GIV promotes lipid-induced macrophage dysfunction and impaired efflux, whereas GIV loss enhances macrophage-driven RCT and protects against atherosclerosis. *Statistics*: All results are presented as mean ± SEM. Significance was assessed using an unpaired t-test or one-way ANOVA. *p*-values < 0.05 were considered significant. *p* < 0.05 (*), *p* < 0.01 (**), *p* < 0.001 (***) and *p* < 0.0001 (****).

Because tissue lipid ‘unloading’ is a defining consequence of enhanced RCT, we quantified fecal sterol excretion by targeted lipidomics as a direct readout of whole-body cholesterol clearance. Myeloid GIV KO mice displayed a robust increase in fecal free sterols and oxysterols, most pronounced under high-fat diet conditions (**Figure 2K, Figure S4,** and **Data S3 and S4**). Notably, this included marked elevations in the injury-inducing oxysterols 7-ketocholesterol (7-KetoC) and 7α-hydroxycholesterol (7α-OHC) (p < 0.0001, **Figure 2L** and **2M**), key mediators of plaque^52–54^ and liver^55^ toxicity. Stool lipidomics also provides direct evidence that loss of myeloid GIV drives systemic cholesterol disposal *via* the fecal route (**Figure 2N**).

Taken together, these findings show that myeloid GIV is a negative regulator of RCT, which promotes lipid accumulation across vascular and metabolic tissues (**Figure 2K**). Its absence unleashes RCT, relieving plaque and organ lipid burden and defining an axis through which macrophages ‘defat’ tissues (**Figure 2N**).

### GIV drives foam cell formation by tilting cholesterol balance toward storage and inflammation

We next asked whether GIV’s ability to inhibit RCT in the setting of systemic lipid overload originates from macrophage-intrinsic differences in cholesterol handling. When challenged with oxLDL, GIV-deficient thioglycolate-elicited peritoneal macrophages (GIV-KO TGPMs, **Figure 3A**) accumulated markedly fewer lipid droplets than WT, as revealed by ORO staining (**Figure 3B** and **3C**). This finding was recapitulated in hyperlipidemic ApoE^⁻/⁻^ backgrounds (AKO vs. DKO), where GIV loss significantly reduced foam cell formation (**Figure 3D-3F**). Similar results were observed in human THP1-derived macrophages (GIV-KO THP1, **Figure 3G**), where GIV depletion led to decreased BODIPY⁺ neutral lipid droplets (**Figure 3H** and **3I**), indicating a conserved role for GIV in regulating lipid storage.

**Figure 3:**
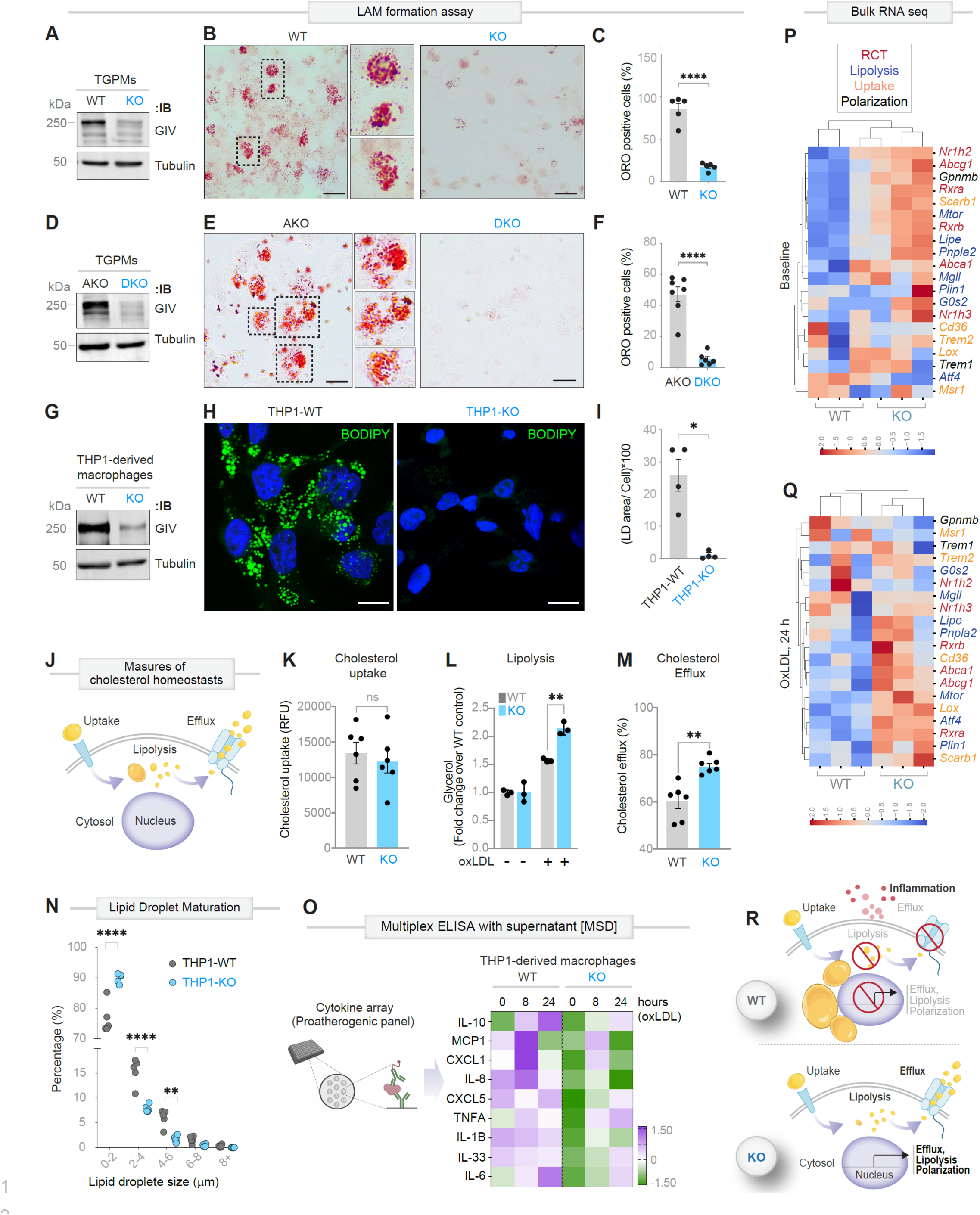
Loss of GIV rewires macrophage cholesterol handling to favor efflux over storage and inflammation. **A-C.** Lipid accumulation in primary macrophages: Immunoblot (A) analysis of thioglycolate-elicited peritoneal macrophages (TGPMs) from WT and GIV-deficient (KO) mice. Representative Oil Red O (ORO) staining (B) of TGPMs stimulated with oxLDL (100 µg/ml, 24 h). Quantification of ORO+ cells shown in (C) (n=5). **D-F**. Genetic validation in a hyperlipidemic background: Immunoblot (D) of TGPMs from ApoE^⁻/⁻^ (AKO) and ApoE^⁻/⁻^;GIV-KO (DKO) mice. Representative ORO staining (E) and quantification (F; n=6). **G-I**. Human macrophage model: Immunoblot (G) of WT vs KO THP1-derived macrophages. Representative BODIPY staining following oxLDL stimulation (100 µg/ml, 24 h; H) with lipid droplet (LD) area quantified per cell (I; n=4). Scale bar, 10 µm. **J-M**. Functional dissection of cholesterol trafficking: Schematic (J) of cholesterol uptake, lipolysis, and efflux steps. Quantification of uptake (K; n=6), lipolysis (L; n=3), and efflux (M; n=6) reveals a shift toward enhanced efflux in GIV-KO macrophages. See also **Figure S4** showing cytotoxicity kinetics were comparable between WT and KO macrophages, indicating that differences in lipid handling are not attributable to altered cell survival under hyperlipidemic conditions. **N.** Altered LD architecture: Frequency distribution of LD sizes highlights reduced LD growth in GIV-KO cells; n=5. **O.** Inflammatory state: Heatmap of secreted cytokines (multiplex ELISA) at 0, 8, and 24 h post-oxLDL stimulation shows broad suppression of pro-atherogenic inflammatory mediators in GIV-KO macrophages (n=3). **P-Q**. Transcriptomic reprogramming: heatmaps in resting (P) and oxLDL-challenged macrophages (Q) show that GIV deficiency downregulates pathways for lipid uptake/storage and inflammatory polarization while upregulating cholesterol efflux signatures. **R**. Model: Working model depicting how GIV promotes foam-cell formation by driving cholesterol uptake, storage, and inflammation while suppressing lipolysis and efflux; GIV loss rebalances these processes toward an atheroprotective state. *Statistics*: All data are mean ± SEM. Significance was assessed using unpaired t-test. p-values < 0.05 were considered significant. *p* < 0.05 (*), *p* < 0.01 (**) and *p* < 0.0001 (****).

To pinpoint affected steps in cholesterol homeostasis, we systematically quantified uptake, lipolysis, and efflux (**Figure 3J**). GIV-KO macrophages displayed unchanged uptake (**Figure 3K**) but exhibited enhanced lipolysis (**Figure 3L**) and increased efflux to extracellular acceptors (**Figure 3M**), signifying a shift from storage toward clearance. Moreover, lipid droplet (LD) size distribution skewed toward smaller droplets in KO macrophages (**Figure 3N**), consistent with defective LD growth, fusion, or storage capacity.

Functionally, cytokine profiling revealed that GIV depletion blunted oxLDL-induced proinflammatory secretions (**Figure 3O**) in TGPMs, highlighting a second inflammation-modulating dimension of GIV in LAMs, coupling lipid storage to inflammatory polarization. Transcriptomic profiling validated these observations, showing broad downregulation of lipid uptake and inflammatory pathways, alongside upregulation of efflux and catabolic programs in KO macrophages, both at baseline (**Figure 3P**) and following oxLDL challenge (**Figure 3Q**). Notably, macrophage survival under lipid-rich conditions was comparable between genotypes (**Figure S5**).

Together, these findings identify GIV as an atherogenic driver that is essential for macrophage foam cell identity, linking cholesterol storage to inflammatory activation (**Figure 3R**). Its loss rewires macrophage metabolism toward lipid efflux, lipolysis, and anti-inflammatory reprogramming, shifting macrophages from atherogenic to atheroprotective, efflux-competent states, without impacting macrophage survival (**Figure 3R**).

### Opposing macrophage programs capture risk and resilience in atherosclerosis

GIV’s role in switching macrophages between atherogenic (when GIV is present) and atheroprotective (when GIV is absent) states (**Figure 3**), prompted us to define the transcriptional programs that distinguish risk from resilience. The human ‘aggravator tLAM’ signature, previously linked to unstable plaques (**Figure 1C**), was robustly induced by oxLDL in WT TGPMs (**Figure 4A**) but blunted in GIV-KO cells (**Figure S6A-B**). Conversely, a 332-gene plaque-stabilization program that is upregulated in human stable plaques was selectively enriched in GIV-KO macrophages after oxLDL challenge (**Figure 4B, Figure S6C**), revealing that GIV restrains a reparative macrophage phenotype.

**Figure 4:**
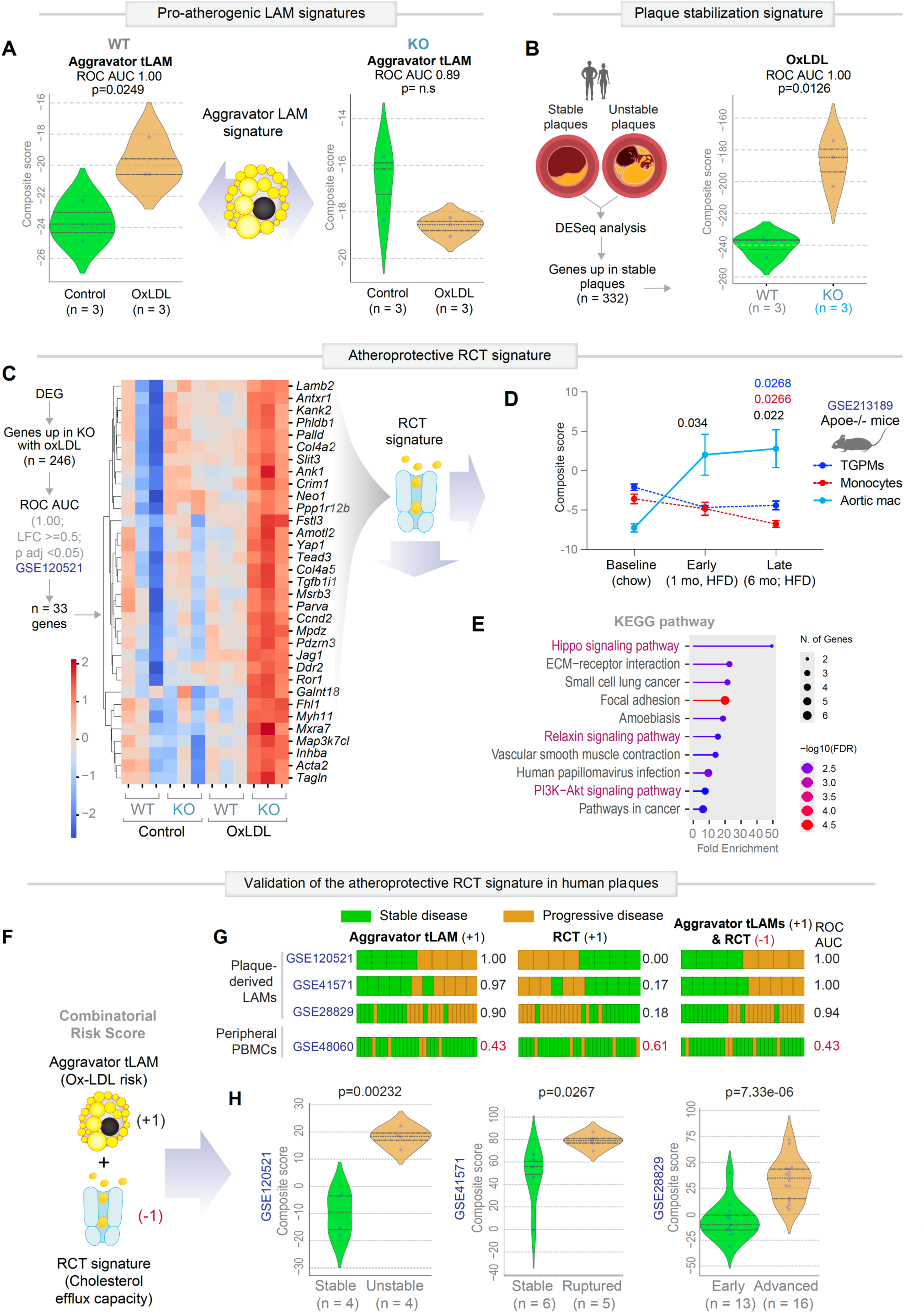
Opposing macrophage gene programs define plaque progression vs. stabilization. **A.** Pro-atherogenic aggravator LAM program: Composite scores for aggravator LAM signature (from **1C**) in WT and KO TGPMs ± oxLDL stimulation. ROC AUC values and p-values indicate classification accuracy. **B.** Human plaque stabilization signature: *<u>Left</u>* - DESeq analysis identifies a plaque-stabilization signature (n = 332 genes, enlisted in **Data S5**) upregulated in human stable plaques. *<u>Right</u>* - Violin plots of stabilization score in WT and KO TGPMs. **C.** Atheroprotective RCT signature: Genes upregulated in GIV-KO macrophages exposed to oxLDL were trained on human plaque transcriptomes ([GSE120521](https://www.ncbi.nlm.nih.gov/geo/query/acc.cgi?acc=GSE83687)^120^; AUC = 1.0, *p* < 0.05) to generate an atheroprotective RCT signature (n=33 genes, enlisted in **Data S6**). Heatmap shows expression of the RCT signature in WT and KO macrophages ± oxLDL. **D.** Validation in diet- and genetic-driven atherosclerosis: Composite RCT scores plotted across macrophage subsets (TGPMs, circulating monocytes, aortic macrophages) in a hyperlipidemic Apoe^-/-^ model (GSE213189^121^) at baseline, 1 mo (early), and 6 mo (late) after exposure to HFD. Black p-values denote comparisons of aortic macrophages relative to baseline across disease stages. Blue and red p-values indicate comparisons of aortic macrophages versus TGPMs and monocytes, respectively, at the indicated time points. **E.** Pathway enrichment: Kyoto Encyclopedia of Genes and Genomes (KEGG) pathway analysis of the RCT signature reveals enrichment for cholesterol efflux and atheroprotective signaling pathways. **F-H**. Integrated clinical risk scoring: Schematic overview (F) of a combined risk score leveraging opposing aggravator LAM (injury-driven progression) and RCT (efflux-driven stabilization) programs. Bar plots (G) show distribution of stable (green) and progressive (beige) lesions ranked by aggravator LAM-only (left), RCT-only (middle), or the integrated (aggravator LAM + RCT; right) score. ROC AUC values report classification accuracy across datasets. Violin plots (H) show performance of the integrated score across human plaques (stable vs. unstable) and mouse lesions (early vs. advanced). *Statistics*: All results are presented as mean ± SEM. Significance was assessed using Welch’s t-test. p-values < 0.05 were considered significant.

Training GIV-KO–induced genes on human plaque transcriptomes distilled an atheroprotective 33-gene RCT signature that perfectly classified disease outcomes (AUC = 1.0, p < 0.05) (**Figure 4C, Figure S6**). This signature was repressed in WT but amplified in KO macrophages under lipid stress, consistent with a GIV-dependent block on cholesterol efflux and resolution (**Figure 4C**). We tested the physiologic relevance of this RCT signature by mapping it across macrophage subsets in a hyperlipidemic Apoe^⁻/⁻^ model (GSE213189). Among circulating monocytes, TGPMs, and aortic macrophages, only the latter exhibited a sustained elevation in RCT signature during HFD exposure, persisting into advanced lesions (**Figure 4D**, cyan solid line). TGPMs displayed transient suppression followed by recovery (**Figure 4D**, blue dotted line), whereas monocytes showed a progressive decline (**Figure 4D**, red dotted line). Pathway enrichment analysis confirmed RCT-associated networks governing cholesterol efflux and anti-inflammatory reprogramming (**Figure 4E**).

Integrating aggravator tLAM (injury-driven; 20-gene) and protective RCT (efflux-driven; 33-gene) programs yielded a combinatorial risk score (**Figure 4F**) that outperformed either signature alone. This combined metric robustly distinguished stable from progressive disease across cohorts (**Figure 4G**) and segregated human stable vs. unstable plaques and early vs. advanced lesions (**Figure 4H**).

Together, these data establish GIV as a molecular checkpoint linking macrophage transcriptional state to clinical plaque behavior, driving vulnerability when present and resilience when removed. The discriminative accuracy of murine digital signatures in human datasets also demonstrates the direct translational relevance of this macrophage-intrinsic switch from murine models to human atherosclerosis.

### GIV tethers ABCA1 to Gαi and LXR and mislocalizes the transporter into endomembranes

To define how GIV suppresses RCT at the protein network level, we focused on the rate-limiting efflux transporter ABCA1. Using *in situ* proximity ligation assays (PLA)^56^ (**Figure 5A**), we visualized endogenous full-length assemblies of **ABCA1●GIV** complexes in single macrophages. These complexes were abundant in WT cells but, as expected, virtually absent in GIV-KO macrophages (**Figure 5B-5D**). Time-course studies revealed that **ABCA1●GIV** complexes are constitutively present at baseline but markedly diminished in fully established LAMs by 24 h after oxLDL exposure (**Figure S7A-C**). Sequential dual-color PLA demonstrated that GIV nucleates ternary **ABCA1●GIV●Gαi** complexes in a ligand-dependent manner, peaking within 1 h of oxLDL stimulation and dissipating by 24 h (**Figure 5E and 5F; S7D-E**). This transient assembly is consistent with GIV’s established role as a dynamic signaling scaffold across diverse receptor systems^57–65^, positioning GIV as a temporal gatekeeper that couples ABCA1 to inhibitory Gαi signaling during early lipid stress. Accordingly, **ABCA1●Gαi** interactions were markedly reduced in GIV-KO cells (**Figure 5G–5I**), confirming GIV’s role as the molecular bridge between the transporter and inhibitory Gαi signaling, as described in other receptor systems^65^.

**Figure 5:**
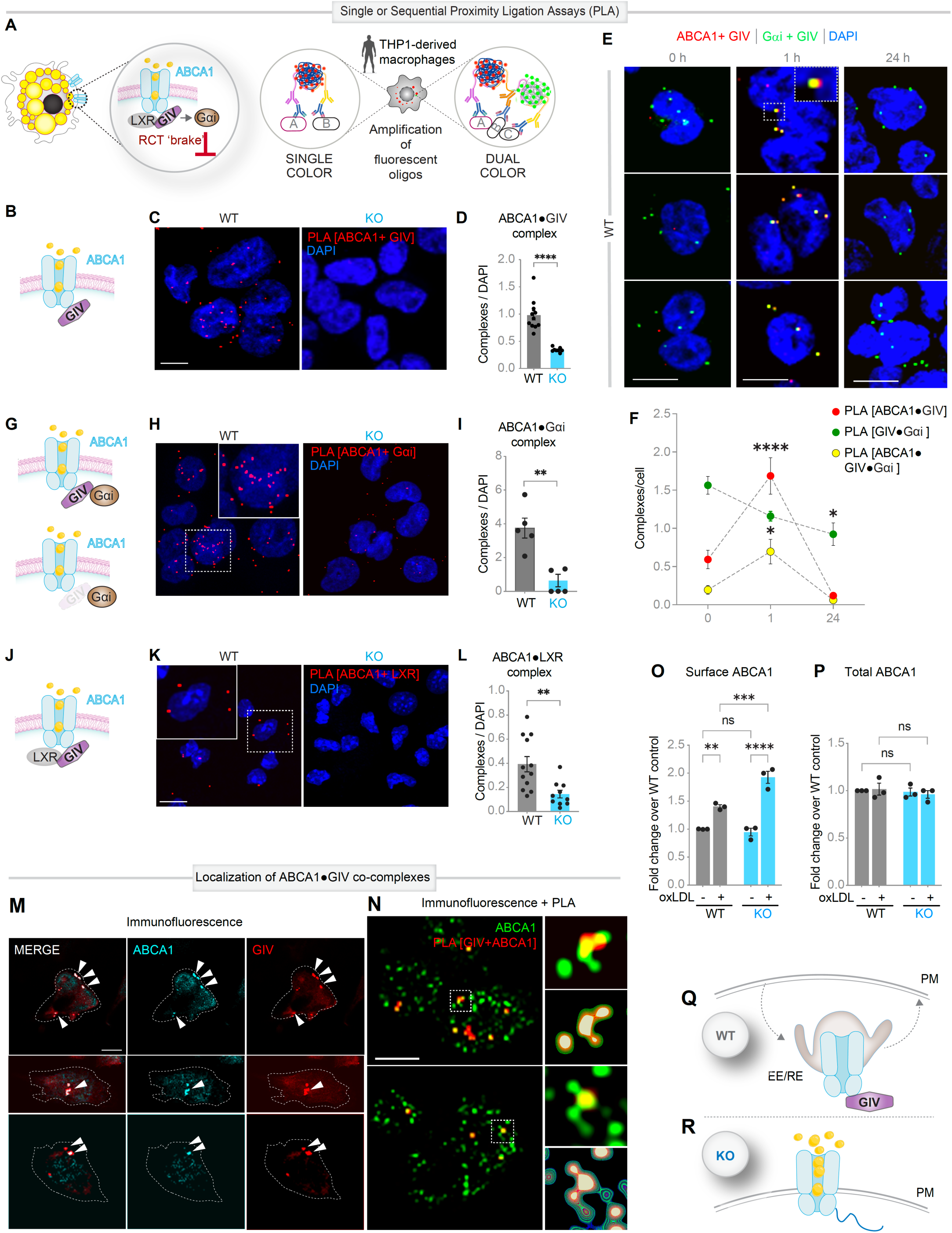
GIV scaffolds ABCA1 to Gαi and LXR, sequestering transporter in endomembranes. **A.** Workflow for *in situ* proximity ligation assays (PLA), implemented either as single- or sequential dual-color, to visualize and quantify ABCA1-bound complexes (B-L) in WT and GIV-KO THP1-derived macrophages. **B-L**. ABCA1●GIV complexes (B) detected by single-color PLA. Representative images (C) and quantification (D). Scale bar, 10 µm. See **Figure S7B and S7C** for dynamics of complexes upon oxLDL (100 µg/ml) treatment for 24h. **E-F**. ABCA1●GIV●Gαi ternary complexes (E) detected by dual-color PLA. Representative images (E) and quantification (F). Scale bar, 10 µm. See **Figure S7D and S7E** for dynamics of complexes upon oxLDL (100 µg/ml) treatment for 1 h and 24h. **G-I**. ABCA1●Gαi complexes (G) detected by single-color PLA. Representative images (H) and quantification (I). **J-L**. ABCA1●LXR complexes (J) detected by single-color PLA. Representative fields (K) and quantification (L). **M**. Representative montage of WT THP1-derived macrophages showing subcellular co-localization of ABCA1 (cyan) and GIV (red) (M). See **Figure S7F** for a montage of additional images showing intracellular sequestration of ABCA1 in lipid-laden WT THP1 macrophages. **N**. Representative montage showing co-localization of ABCA1 (green) with ABCA1●GIV complexes (visualized by single color PLA, red). Insets highlight ABCA1-positive endosomal compartments (green) with co-complex–positive microdomains (red). **O-P**. Surface (O) and total ABCA1 (P) were measured by flow cytometry and quantified using NovoExpress software. See **Figure S7G** for surface ABCG1. **Q-R**. Working model illustrating how GIV sequesters the ABCA1 transporter within endomembranes; in the absence of GIV, ABCA1 remains localized to the plasma membrane, enabling cholesterol efflux. RE, Recycling Endosomes; EE: Early Endosomes *Statistics*: All results are presented as mean ± SEM. Significance was assessed using an unpaired t-test or one-way ANOVA. *p*-values < 0.05 were considered significant. *p* < 0.01 (**), *p* < 0.001 (***) and *p* < 0.0001 (****).

GIV also influenced ABCA1’s regulatory hierarchy: in WT macrophages, GIV shifted ABCA1’s binding preference toward its binding partner LXR, enhancing the formation of **ABCA1●LXR** complexes (**Figure 5J–5L**), which signals a state when both are functionally inert^66^; this interaction was diminished in GIV-KO cells. Beyond reshaping protein partnerships, GIV altered ABCA1’s spatial topology. Confocal imaging showed that ABCA1 and GIV co-localize within lipid-laden LAMs (but not within lipid droplets; **Figure 5M; Figure S7F**, where ABCA1-positive endomembranes were enriched in GIV-containing subdomains. At higher resolution, the ABCA1●GIV co-complexes in WT cells appeared to be concentrated in ABCA1-positive intracellular compartments that frequently displayed tubular extensions (**Figure 5N**), with a concomitant suppression of their abundance at the cell surface (**Figure 5O** and **5P**); this suggests that GIV traps the transporter in either sorting (early) or recycling endosomal microdomains. Of note, GIV’s influence on cell-surface localization of RCT-enabling transporters is selective for ABCA1 and does not extend to its paralog ABCG1 (**Figure S7G**), underscoring a mechanistic specificity in how GIV controls cholesterol efflux.

Collectively, these results identify GIV as a scaffolding hub that tethers ABCA1 to Gαi and LXR and sequesters the inert transporter in endomembranes (**Figure 5Q** and **5R**). It raises the possibility that coupling of ABCA1 to Gαi might suppress cAMP→PKA→CREB signaling and that anchoring LXR might curtail transcriptional support for efflux genes; both mechanisms converge to enforce an efflux-suppressive state to favor LAM genesis.

### GIV’s C-terminus binds Ser^2054^, the constitutive efflux-activation switch on ABCA1

We next asked how GIV’s scaffolding of ABCA1 suppresses cholesterol efflux. Co-immunoprecipitation of endogenous complexes (**Figure 6A**) corroborated PLA data (**Figure 5**), confirming that GIV constitutively associates with ABCA1 and stabilizes an LXR-bound, inert transporter state^66^. In GIV-KO macrophages, LXR association with ABCA1 was markedly reduced despite elevated LXR levels (**Figure 6A**), consistent with a transcriptionally ‘poised’ efflux-competent baseline state (**Figure 3P**).

**Figure 6:**
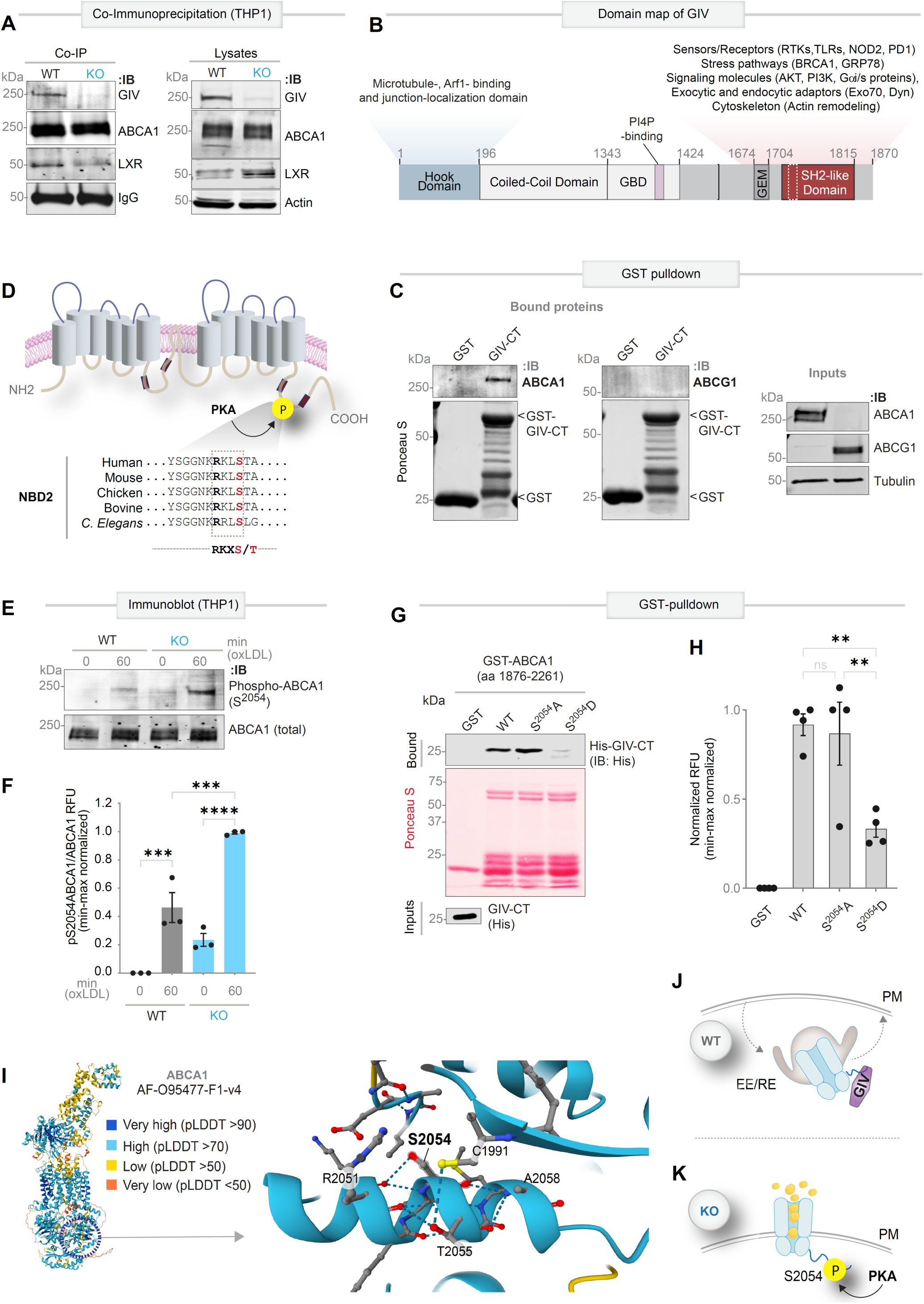
GIV’s C-terminus binds and interferes with the Ser^2054^ efflux-activation switch on ABCA1. **A.** Co-immunoprecipitation (IP) of endogenous ABCA1–bound complexes in THP1-derived macrophages. Lysates (Input) and IP fractions were probed for GIV, LXR, ABCA1, and loading controls (IgG, actin). **B.** Domain organization of GIV showing its C-terminal effector module, which connects to diverse intracellular and cell-surface localized sensors and/or receptors. GBD, Gα-binding domain; GEM, guanine-nucleotide exchange modulator; SH2, Src-like homology-2. **C.** GST pulldown assays using bead-immobilized GST–GIV-CT and lysates of HEK293T cells exogenously expressing untagged, full-length ABCA1 or ABCG1, probed for transporter binding (see **Figure S7** for additional validation using untagged and FLAG-tagged full-length ABCA1 constructs). **D.** Topology map of ABCA1 highlighting the critical PKA site Ser^2054^ within NBD2, a known regulator of efflux function. function. **E-F**. Immunoblot of phospho-ABCA1 (Ser^2054^) in THP1-derived macrophages at baseline and following oxLDL stimulation. Quantification of phospho-ABCA1 normalized to total ABCA1 from three independent biological replicates is shown in F. **G-H**. GST pulldown assays (G) using soluble His–GIV-CT (aa 1660-1870) and bead-immobilized, GST-tagged wild-type (WT), non-phosphorylatable (S^2054^A), or phosphomimic (S^2054^D) ABCA1 NBD2-CT fragments (aa 1876-2261). GIV binding is selectively lost with the S^2054^D mutant. Quantification of four independent biological replicates is shown in H, n=4. **I**. Alphafold structural predictions [AF-O95477-F1-v4; 7 structures in PDB] highlight H-bonds formed by Ser^2054^ contributing key H-bond interactions within a short α-helix. High confidence pLDDT (predicted Local Distance Difference Test) scores support model fidelity. **J-K**. Working Model: In WT macrophages (J), GIV’s C-terminus binds the Ser^2054^ region of ABCA1, retaining the transporter in phospho-suppressed endomembranes and favoring LAM formation. In GIV-KO cells (K), phosphoactivated ABCA1 remains at the plasma membrane (PM), driving cholesterol efflux and restoring RCT. RE, recycling endosomes, EE, early endosomes. *Statistics*: All results are presented as mean ± SEM. Significance was assessed using one-way ANOVA. *p*-values < 0.05 were considered significant. *p* < 0.01 (**).

GIV is a large, multi-modular scaffold (1870 amino acids), whose ∼210 amino acid C-terminal (CT) module contains short linear motifs (SLIMs) that bind diverse cell surface proteins (**Figure 6B**). Biochemical mapping identified this CT module as sufficient for ABCA1 interaction. GST pulldown assays demonstrated selective binding of GIV-CT to ABCA1, but not its paralog ABCG1, confirming transporter selectivity (**Figure 6C**; **Figure S8**). Notably, ABCA1, but not ABCG1, harbors Ser^2054^ within its nucleotide-binding domain 2 (NBD2), a constitutive PKA-targeted phospho-switch that enhances efflux without altering ApoA1 binding or transporter stability^14^ (**Figure 6D**). We therefore asked whether GIV suppresses efflux by engaging this site. Upon oxLDL challenge, WT macrophages displayed minimal PKA-dependent phosphorylation at Ser^2054^, whereas GIV-deficient cells showed ∼2.5-fold increase in phosphorylation (**Figure 6E and 6F**). GIV bound wild-type (WT) ABCA1 and the non-phosphorylatable S^2054^A (Alanine) mutant, but not the phosphomimetic S^2054^D (Aspartate) variant (**Figure 6G, H**), suggesting that phosphorylation at S^2054^ may disrupt GIV•ABCA1 complexes.

Structural modeling provided a mechanistic explanation. Alphafold predictions showed that Ser^2054^ forms stabilizing H-bonds within a short α-helical stretch (**Figure 6I**), suggesting that phosphorylation induces conformational changes that favor efflux activation. By engaging ABCA1 at or near Ser^2054^, GIV likely clamps NBD2 in a non-phosphorylated, inactive conformation, which is released upon PKA-mediated phosphorylation to permit efflux.

Together with findings in **Figure 5**, these results identify Ser^2054^ as GIV-targeted efflux control node (**Figure 6J** and **6K**) and support a model that GIV sequesters inert ABCA1 in endomembranes, likely recycling and/or early endosomes, curtailing phosphoactivatable efflux and promoting LAM formation. In contrast, loss of GIV permits phosphoactivated ABCA1 localization at the plasma membrane, sustained cholesterol efflux, and restoration of RCT.

### GIV●Gαi control of the cAMP–PKA–CREB axis gates ABCA1 activity and enforces LAM fate

We next dissected the upstream signaling circuitry that controls the functional and physical interactions of GIV and ABCA1. In macrophages, GIV is expected to activate Gαi, which in turn should reduce cAMP and blunt PKA and CREB activities, thereby diminishing functional and transcriptional activation of ABCA1 (**Figure 7A**). Consistent with this mechanism, GIV deletion restored intracellular cAMP (**Figure 7B**), enhanced PKA activity (**Figure 7C, Figure S9A**) and CREB-driven transcription (**Figure 7D** and **7E**) following oxLDL challenge. Notably, this elevation in cAMP signaling was not restricted to in vitro systems; cAMP levels were also significantly increased in aortic root macrophage-rich regions from DKO mice compared to AKO controls (**Figure S9B-C**), supporting the in vivo relevance of this GIV●Gαi–cAMP axis in atherogenic niches.

**Figure 7:**
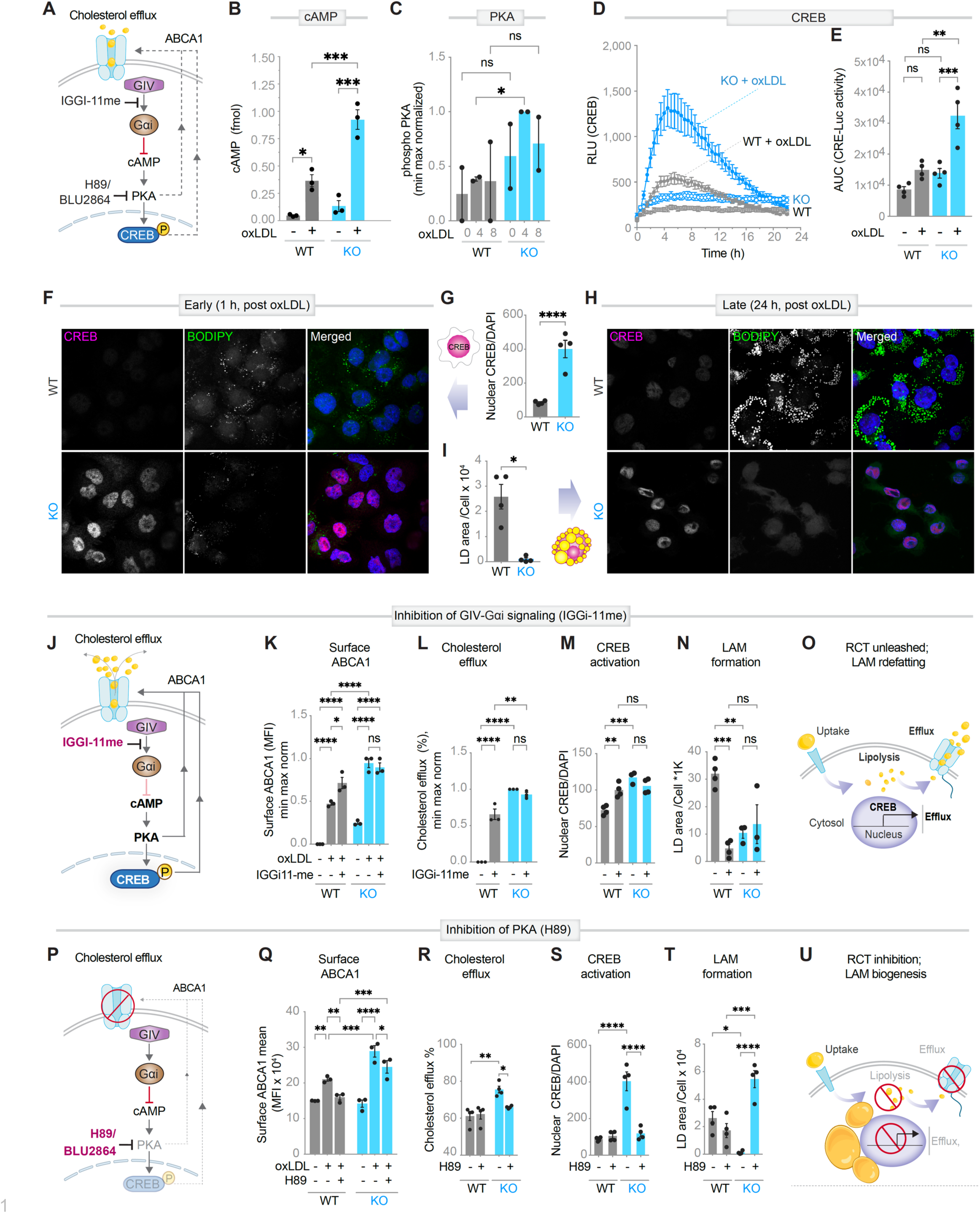
GIV●Gαi control of the cAMP–PKA-CREB axis gates ABCA1 activity and enforces LAM fate. **A.** Mechanistic schematic illustrating an integrated pharmacogenomic approach to dissect how GIV activates Gαi to suppress cAMP–PKA-CREB signaling, thereby reducing phosphoactivation and transcriptional activation of ABCA1. This dual inhibition (‘brake’) ABCA1-mediated cholesterol efflux^9^ and CREB-dependent transcription^15^. IGGi-11me disrupts GIV●Gαi coupling; H89 and BLU2864 inhibit PKA downstream. **B.** cAMP levels in WT vs. GIV-KO THP1-derived macrophages ± oxLDL; n=3. **C-E**. PKA and CREB activation: Quantification of phospho(p)-PKA in WT vs. GIV-KO macrophages (C; n=2, see immunoblots in **Figure S9A**. CRE-Luc activity (D) and corresponding AUC (E) in WT vs. GIV-KO macrophages demonstrating sustained CREB activation in KO cells following oxLDL; n=4. **F-I**. Nuclear CREB and lipid droplet dynamics: Immunofluorescence micrographs and quantification of lipid droplets (BODIPY, green) and nuclear CREB (red; DAPI, blue) at early (1 h, F and G) and late (24 h, H and I) time points after oxLDL exposure. See **Figure S9D-H** for full time-course analysis. **J-O**. GIV●Gαi inhibition favors defatting: Schematic of IGGi-11me intervention (10µM, J), and its impact on surface ABCA1 (K; n=3), cholesterol efflux (L; n=3), CREB activation (M; n=4), and LAM burden (N; n=3). Model (O) summarizing how GIV●Gαi inhibition (10 µM) relieves efflux constraints and drives LAM defatting. **P-U**. PKA inhibition enforces foam cell fate: Schematic of H89 intervention (30µM, P), and its effects on surface ABCA1 (Q; n=3), cholesterol efflux (R; n=4), CREB activation (S; n=4), and LAM burden (T; n=4). Model summarizing how PKA inhibition promotes RCT inhibition and LAM biogenesis (U). See also **Figure S9I-K** for supporting data using an alternative highly potent and specific small molecule (BLU2864, 10 µM)^122^. *Statistics*: All results are presented as mean ± SEM. Significance was assessed using an unpaired t-test or one-way ANOVA. *p*-values < 0.05 were considered significant. *p* < 0.05 (*), *p* < 0.01 (**), *p* < 0.001 (***) and *p* < 0.0001 (****).

We linked these signaling changes to LAM biogenesis using multiplexed single-cell analyses that simultaneously tracked nuclear CREB localization and lipid accumulation (**Figure S9D**). GIV-KO macrophages exposed to oxLDL formed fewer lipid droplets yet showed pronounced nuclear CREB enrichment compared to WT controls, both at early and late stages of LAM formation (**Figure 7F-7I**). Thus, loss of GIV uncouples oxLDL injury from the LAM trajectory, redirecting macrophages toward an efflux-competent state.

To test whether GIV-dependent coupling to Gαi is mechanistically sufficient to restrain cholesterol efflux, we used IGGi-11me^67^, a previously characterized inhibitor that selectively disrupts GIV•Gαi interaction. Disrupting GIV●Gαi coupling with IGGi-11me (**Figure 7J**) restored surface ABCA1 (**Figure 7K**), enhanced cholesterol efflux (**Figure 7L**), reactivated CREB (**Figure 7M**), and suppressed LAM formation (**Figure 7N, Figure S9E and S9F**), thereby unleashing RCT and driving LAM ‘defatting’ (**Figure 7O**). Conversely, direct PKA inhibition with H89 (**Figure 7P**) impaired ABCA1 function (and **Figure 7Q**), reduced efflux (**Figure 7R**), dampened CREB activity (**Figure 7S**), and exacerbated LAM formation (**Figure 7T; Figure S9G** and **S9H**), thereby inhibiting RCT and driving LAM biogenesis (**Figure 7U**). These effects were recapitulated with the highly selective PKA inhibitor BLU2864^68^ (**Figure S9I-K**).

Collectively, these findings define a GIV●Gαi–cAMP checkpoint that gates the cAMP–PKA–CREB axis, suppresses ABCA1 activity, and locks macrophages in a pro-LAM, efflux-refractory state, revealing a druggable node for reprogramming macrophage fate and restoring RCT.

### The GIV●Gαi–cAMP checkpoint represents a therapeutic entry point for RCT restoration

To test the therapeutic potential of targeting the GIV●Gαi–cAMP checkpoint, we quantified LAM formation *in vivo* using an established model^69^, i.e., macrophages recovered from diet-induced hyperlipidemic mice, with or without pharmacological interventions (**Figure 8A**). Flow cytometric (**Figure 8B** and **8C**) and fluorescence image (**Figure S10A**) analysis of BODIPY-labeled neutral lipids revealed a significant reduction in LAM burden in GIV-KO macrophages compared to WT controls, a phenotype that was phenocopied by pharmacologic disruption of GIV●Gαi signaling (IGGi-11me). These findings indicate that compartment-specific disruption of GIV●Gαi signaling in peritoneal macrophages is sufficient to suppress foam-cell formation *in vivo*.

**Figure 8:**
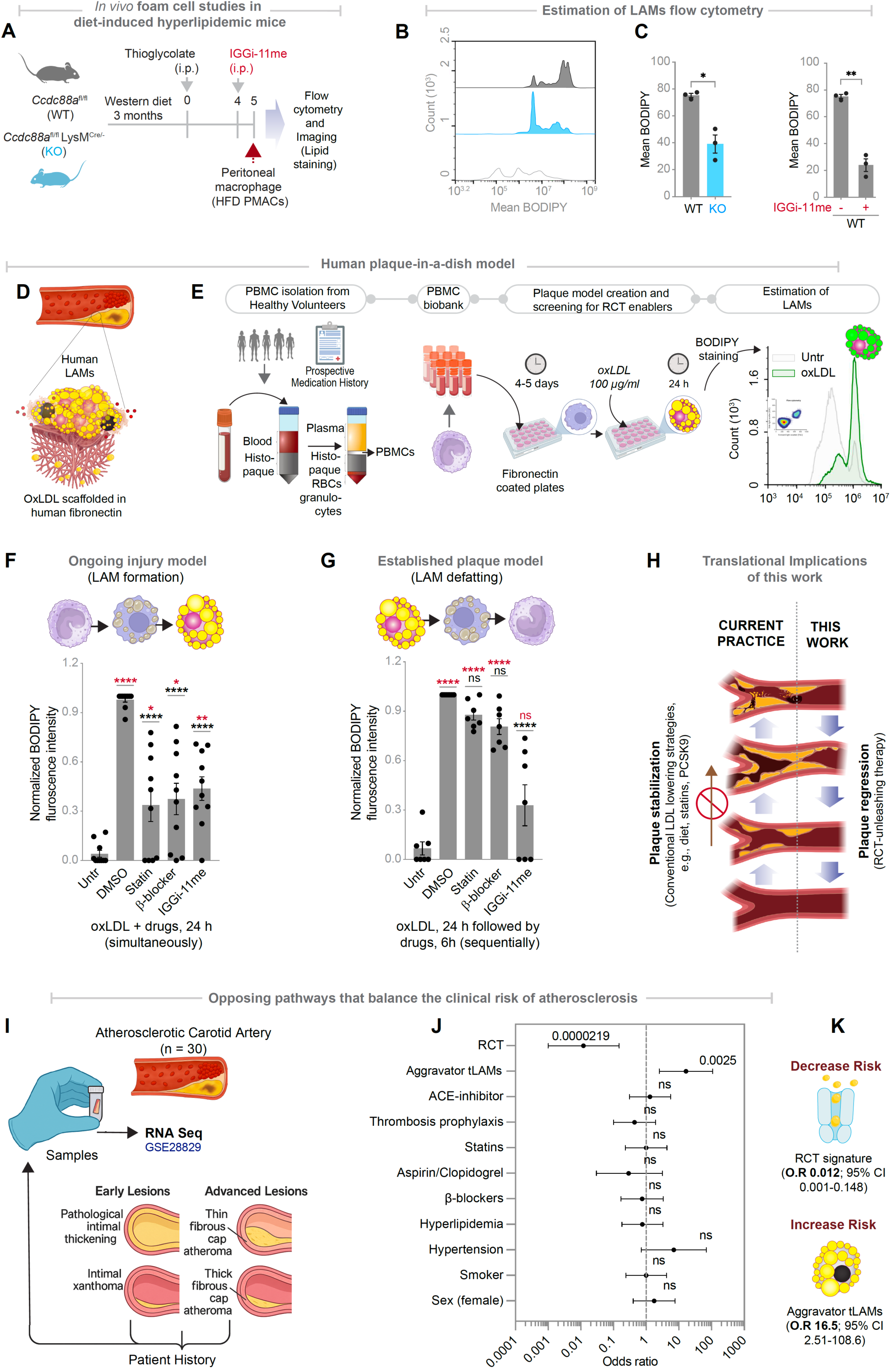
Therapeutic modelling reveals the clinical impact of RCT restoration. **A.** Study design for *in vivo* disruption of the GIV●Gαi interface in diet-induced hyperlipidemic mice. Eight-week-old mice were fed a high-fat diet (HFD) for 3 months, followed by thioglycolate elicitation and acute pharmacologic inhibition of GIV●Gαi signaling using intraperitoneal (i.p, 1 mg/kg) IGGi-11me. Peritoneal macrophages (HFD-PMACs) were isolated and analyzed for lipid burden by flow cytometry and imaging (n=3). **B-C**. Representative BODIPY histograms (B) and quantification of neutral lipid accumulation (C), as determined by flow cytometry. See **Figure S10A** for orthogonal analyses of lipid burden by confocal imaging. **D**. Schematic of a simple, scalable human plaque-in-a-dish model in which oxLDL transforms PBMC-derived macrophages into LAMs scaffolded on human fibronectin. **E-G**. Workflow (E) for therapeutic testing in human PBMC-derived macrophages. Compounds were administered either concurrently with oxLDL to assess blockade of LAM formation (F; n=10) or after LAM establishment to assess reversal (“defatting”) capacity (G; n=7). Bar graphs compare the efficacy of statin (Atorvastatin, 10 µM), β-blocker (Metoprolol, 10 µM), and IGGi-11me (10 µM) under each condition. See also **Figure S10B-D** for impaired reversal in statin-experienced macrophages. **H**. Conceptual framework contrasting current standard-of-care LDL-lowering strategies (left) with RCT-restoring approaches enabled by targeting the GIV–ABCA1 axis (right; this work). **I-J**. Study design (K) and forest plot (L) depicting *in silico* projection of pro-atherogenic aggravator tLAM vs anti-atherogenic RCT risk signatures in an independent cohort of patients with carotid atherosclerosis. **K**. Model. Opposing lipid pathways define clinical outcomes: oxidized lipid burden and aggravator LAM states drive disease progression, whereas restoration of RCT promotes resilience and reduces atherosclerotic risk. See also **Table S2** for clinical correlates and **Data S7** for odds ratio calculations. **Figure S10E and Table S3** summarize lipid-linked risk modifiers validated across Phase III clinical studies. *Statistics*: Data shown as mean ± SEM. Significance was assessed by unpaired t-test or one-way ANOVA. In panels C and D, red p-values denote comparisons to untreated controls (Untr), black to oxLDL + DMSO controls. Significance: p < 0.05 (), < 0.01 (), < 0.001 (), < 0.0001 (****).

To extend these observations to human systems, we engineered a simple, scalable human plaque-in-a-dish model, in which PBMC-derived macrophages were scaffolded on human fibronectin and transformed into LAMs upon oxLDL exposure (**Figure 8D and 8E**). Fibronectin was chosen because it is the key extracellular-matrix component deposited under atheroprone hemodynamics^70^, efficiently scaffolding oxLDL to recruit monocytes and vascular smooth-muscle cells (VSMCs)^71^. This microenvironment embodies the classical *response-to-retention* paradigm^72^, where trapped lipoproteins initiate monocyte invasion, rapid LAM differentiation, and inflammatory amplification within plaques destined to progress^72^.

We next asked whether pharmacologic interventions could block or reverse LAM formation. In co-treatment assays, where drugs and oxLDL were applied concurrently, statins and β-blockers effectively blocked LAM formation, as expected (**Figure 8F**). This confirmed that the system faithfully recapitulates LAM biogenesis and provides a tractable platform for therapeutic testing. IGGi-11me, which disrupts GIV●Gαi coupling, also markedly suppressed LAM formation (**Figure 8F**).

In reversal assays, whereas LAMs were first established before treatment, statins and β-blockers failed to defat LAMs, while IGGi-11me restored an efflux-competent phenotype, lipid-poor phenotype (**Figure 8G**). Consistent with clinical observations that chronic statin use downregulates ABCA1^73,74^, IGGi-11me was ineffective in PBMCs derived from statin-treated donors (**Figure S10B-D**).

Together, these results standardize a scalable, simple human new approach methodology (NAM) that can be used for RCT-enabler drug screening and identify GIV●Gαi inhibition as a druggable checkpoint in human LAM biology, capable of restoring RCT where conventional therapies fail (**Figure 8H**). While IGGi-11me was effective *in vivo* in mice when assessing peritoneal LAMs (**Figure 8C**), its undetectable systemic concentrations in plasma^67^ precluded further evaluation of systemic therapeutic efficacy across organ systems.

### LAM-derived molecular programs mirror clinical lipid risk axes

Circulating lipid levels remain the cornerstone of cardiovascular risk assessment. For example, elevated LDL and oxidized phospholipids^75–80^ correlate with heightened risk, whereas HDL levels^75,76,79^ and cholesterol efflux capacity (CEC)^77–79^ are protective (**Figure S10E**). In clinical practice, ratios of these opposing measures, i.e., LDL/HDL or oxLDL/CEC remain gold standards for risk stratification^81^. We next asked whether transcriptional programs of LAM biology could serve as molecular surrogates for these clinical traits.

We tested whether the pro-atherogenic aggravator LAM signature (**Figure 1C**) correlates with high-risk (LDL/oxLDL), and whether the atheroprotective RCT signature (**Figure 4C**) mirrored established protective factors (HDL/CEC).

To quantitatively test this relationship, we integrated both signatures into a combinatorial molecular risk model (**Figure 8I-K**) and applied it to a validation cohort of patients with carotid atherosclerosis (n = 30; **Figure 8I** and **8J**). High aggravator LAM scores correlated with ∼16.5-fold risk of disease progression, whereas high RCT scores conferred >98% protection (OR = 0.012, 95% CI 0.001–0.148; p = 2.19 × 10⁻⁵; **Figure 8K**).

Collectively, these findings demonstrate that restoring RCT through GIV●Gαi inhibition reprograms macrophage fate, defats human plaques ex vivo, and aligns molecularly with the clinical axis of lipid resilience. This framework unifies mechanistic discovery, human modeling, and risk prediction, thereby positioning RCT restoration as a next-generation therapeutic paradigm for immunometabolic disease.

## DISCUSSION

### Macrophage Lipid Persistence as the Unmet Barrier to Regression

Despite decades of lipid-lowering therapy, macrophage lipid persistence remains the central obstacle to atherosclerosis regression. Statins stabilize lesions by lowering LDL but fail to purge lipid reservoirs sequestered in plaque macrophages. Our study identifies GIV (*CCDC88A*) as a macrophage-intrinsic metabolic checkpoint that couples G-protein signaling to RCT. By scaffolding ABCA1 to Gαi and LXR, GIV suppresses the cAMP–PKA–CREB axis, sequesters ABCA1 in endomembranes, and locks macrophages into a lipid-laden, inflammatory LAM state. Genetic loss or chemical disruption of this axis restores ABCA1 phosphorylation and efflux competence, mobilizing hepatic and adipose lipids and reducing plaque burden. Integrating an end-to-end human framework of macrophage-continuum modeling, *ex vivo* plaque-in-a-dish assays, and patient-derived transcriptional networks positions GIV●Gαi–ABCA1 node as a tunable axis of risk versus resilience. These findings reframe macrophages from passive foam cells to programmable effectors of lipid clearance, establishing RCT restoration as a new therapeutic paradigm for immunometabolic disease.

### Opposing Macrophage Programs Govern Plaque Risk and Resilience

Building on the SMaRT macrophage-continuum model, we delineated two opposing macrophage states, alleviator reactive (rLAM) and aggravator tolerant (tLAM), that align with plaque progression and stabilization, respectively. Unlike prior scRNA-seq taxonomies^24,82^ that emphasize cell diversity at the expense of mechanism (e.g., the now-recognized double-edged role of Trem2 in LAMs), this continuum-based approach grounds macrophage biology in disease trajectories.

Aggravator LAMs carried a risk signature for progression, and within this set, we identified *CCDC88A* (encoding GIV) as a brake on RCT, a non-intuitive mechanism that would likely have been obscured by cell-centric clustering. Importantly, the reproducibility of this continuum-based approach to interrogating macrophage states across tissues underscores its strength. For example, in IBD, the discovery of iColAMs and niColAMs revealed that non-inflammatory colitis-associated macrophages deploy NOD2 as a brake on inflammation^57^.

Together, these results support a unifying model: immune homeostasis arises from counterpoised macrophage populations—accelerators that drive inflammation and brakes that resolve it. Pathology ensues when the brakes get stuck (as in atherosclerosis) or fail (as in IBD).

### Macrophage Activation as a Context-Dependent Regulation

Our prior work showed that GIV skews innate immune signaling by selectively dampening proinflammatory cytokine production, while permitting anti-inflammatory IL10 responses downstream of selective TLRs, e.g., TLR4^50^ and the cytosolic sensor, NOD2^57^. Thus, GIV **biases signals** towards macrophage tolerance and induces the pro-atherogenic IL10^83,84^. By contrast, the macrophage responses examined here are triggered by sterile oxidized lipoprotein stress and reflect lipid-driven cellular stress rather than canonical pathogen-sensing pathways. Under these conditions, GIV amplifies oxLDL-induced pro-inflammatory programs, with no discernible impact on anti-inflammatory IL10 responses. GIV acts as an anti-inflammatory regulator during infection yet promotes inflammatory remodeling under lipid overload, illustrating context-dependent deployment of the same molecular checkpoint.

Positioned at the intersection of innate immune sensing and metabolic stress, GIV may function as a signaling hub for receptors such as TLR4^50^ and NOD2^57^, both established drivers of plaque inflammation^85–89^. Murine genetic models have confirmed that both TLR4^85–87^ and NOD2^88,89^ are required for plaque progression and macrophage differentiation into LAMs, processes that may be further potentiated by feedforward ABCA1-PKA signaling^90^. Proposed mechanisms include suppression of NFkB signaling and skewed induction of the pro-atherogenic and pro-survival cytokine IL10^83,84^. Our findings suggest that this skewing toward tolerogenic signaling represents a convergent mechanism by which GIV may link innate immune sensing to impaired lipid clearance and LAM-fate stabilization during atherogenesis.

### The GIV●Gαi Molecular Switch Emerges as a Unified Brake on Cholesterol Efflux

Mechanistically, GIV suppresses cholesterol efflux through a dual mechanism: (i) structural inhibition, and (ii) signal-mediated suppression.

First, GIV binds directly to ABCA1 at or near the critical Ser^2054^ within its NBD2 domain which is a constitutively phosphorylatable PKA site that is essential for efflux function^9^. That phosphorylation of ABCA1 at Ser^2054^ upon oxLDL stimulation is higher in the absence of GIV raises the possibility that GIV may act as a steric competitor^91^, preventing phosphorylation and conformational changes in NBD necessary for transporter activation (such as nucleotide binding/hydrolysis)^9,92^. Because GIV loses binding to the phosphomimetic mutant (S^2054^D) but not S^2054^A suggests the possibility of a feedback-coupled ‘brake’ system in which rising cAMP→PKA signals trigger ABCA1 phosphorylation at Ser^2054^, which releases GIV, toggling the transporter into its active state. Second, GIV couples ABCA1 to Gαi, dampening the cAMP, PKA-, and CREB-pathways that are indispensable for both transporter activity^9^ and transcriptional^15^ induction.

Together, these two layers constitute a potent, evolutionarily conserved molecular “brake” linking cAMP signaling to ABCA1 activity, substantiating cAMP’s widely established role as a potent anti-atherogenic pathway^15–20,93–95^, resolving long-standing uncertainty over how PKA inputs regulate efflux.

### ABCA1 Efflux as a Context-Dependent Toggle

That multiple models of ABCA1 regulation^96^ coexist in the literature underscores how elusive its mechanism has been. These mechanisms, ranging from clathrin-dependent recycling to caveolar lipidation,^96^ may in fact represent context-specific states of its functional toggle. Our data suggest that these models may not be conflicting, but context-dependent, as ABCA1 toggles between GIV-bound (efflux-restrained; brake ‘on’) and GIV-free (efflux-competent; brake ‘off’) conformations. In GIV-rich aggravator LAMs, ABCA1 is sequestered in endomembranes with LXR, consistent with suppressed efflux; in GIV-deficient macrophages, it relocates to the plasma membrane, thereby engaging ATP-driven dimerization and lipid-loading cycles. GIV colocalization with ABCA1 at peripheral discoid patches and perinuclear vesicles dovetails with endocytic recycling models^96–100^ invoking caveolae-associated lipidation, clathrin-dependent endocytosis, and Rab4-dependent endocytic recycling^101^, while GIV depletion aligns with inefficient endocytosis favoring surface cycles of ATP-driven ABCA1 dimerization, lipid loading, and apoA-I engagement. Because total ABCA1 remained similar regardless of GIV, Arf6-dependent ABCA1 degradation^102^ does not appear to be impacted. This model integrates transcriptional and post-translational regulation within a single allosteric framework: phosphorylation at Ser^2054^ serves as the trigger to release GIV, unleash PKA activity, and restore RCT. Such toggle-like control allows macrophages to switch rapidly between lipid-storage and lipid-export states depending on GIV occupancy and cAMP tone. It is also possible that GIV has additional binding sites that offer specificity of interaction with ABCA1 (vs ABCG1).

### Reprogramming of Macrophage Fate and Impact of RCT-enablers

Our findings illuminate why current immunometabolic therapies incompletely resolve plaque lipid burden. Statins, ACE inhibitors^103,104^, β-blockers, and antithrombotic agents, though effective at lowering LDL or stabilizing advanced plaques^105–107^, cannot mobilize the vast lipid reservoirs sequestered within macrophages. β-blockers, which chronically dampen Gαs signaling, stabilize plaques^108^ via immunomodulatory pathways but do not mobilize cholesterol; are believed to attenuate adrenergic stress and mechanical strain on plaque. GIV●Gαi directly governs lipolysis and cholesterol efflux. GIV operates at the complementary Gαi node, suppressing cAMP→PKA →CREB signaling to block efflux, and thus represents a unique therapeutic lever. Inhibiting this checkpoint defats macrophages without systemic lipid lowering or broad immunosuppression, addressing the root of residual cardiovascular risk.

Equally important is how GIV inhibition contrasts with emerging Trem2-based interventions. Trem2 activation expands LAMs and stabilizes advanced plaques by reducing inflammation and plaque necrosis, but it carries a double-edged liability^29,109^: in early lesions, Trem2 drives maladaptive macrophage survival and accelerates plaque growth^28,29,45^. This stage-dependence has tempered enthusiasm for Trem2 as a universal therapeutic target. By contrast, GIV depletion elicits robust macrophage defatting and efflux, stimulates lipolysis, and dampens inflammation without affecting macrophage survival -- features expected to be beneficial across both early and advanced lesions, offering a more uniform therapeutic window. This breadth of efficacy positions GIV●Gαi inhibition as a safer, more versatile approach for reversing immunometabolic pathology.

### Human NAMs Bridge Mechanism and Medicine

Our findings redefine how macrophage-driven lipid retention can be studied and therapeutically targeted. The fibronectin-based human NAM we developed distills the atheroprone niche to its essentials, i.e., matrix retention, monocyte adhesion, and lipid stress, creating a minimal yet mechanistically faithful model of plaque biology. Its simplicity is its strength: scalable, reproducible, and fully aligned with the federally mandated NAM standards for human-relevant testing. This platform transforms macrophage efflux from a descriptive process into a quantifiable, druggable endpoint. By recreating the conditions that drive LAM formation, it enables high-throughput screening of RCT modulators in a system that preserves human variability, e.g., sex, treatment history, and metabolic background.

Integrating these NAMs with cohort-level anchoring using ‘digital biomarkers’ such as aggravator LAM and RCT gene signatures, closes the translational loop: molecular interventions can now be linked to cellular reprogramming and simulated clinical benefit. Together, this mechanism-to-model-to-patient continuum converts RCT restoration from a theoretical concept into a predictive and actionable therapeutic paradigm for immunometabolic disease.

### Integrating Mechanistic and Systems Insight

At a systems level, the GIV●Gαi → cAMP/PKA/CREB inhibitory circuit emerges as a master integrator that controls ABCA1 function, macrophage fate, and tissue-level lipid balance. Our findings support a hierarchical model in which GIV●Gαi-dependent inhibition of cAMP→PKA signaling functions as the primary upstream “brake,” while enhanced PKA/CREB→ABCA1-mediated cholesterol efflux and RCT represent downstream metabolic consequences of releasing that brake. Thus, signaling and metabolic outputs are mechanistically coupled rather than independent drivers of the observed phenotype. This mechanism unites transcriptional, post-translational, and structural regulation into a single, druggable node. Disabling this checkpoint re-engages RCT, reversing the defining cellular pathology that resists current therapies. The phenotype observed across vascular, hepatic, and adipose compartments suggests broad applicability to obesity and fatty-liver disease, where lipid-engorged macrophages drive chronic inflammation.

## Study Limitations

While validated across independent human plaque datasets, extension of the macrophage-continuum model^44^ to other vascular beds remains to be tested. Although macrophages are central to atherosclerotic plaque development, other immune cell types, including lymphocytes, also contribute to disease progression^110,111^; thus, the impact of myeloid GIV deficiency on broader plaque immune composition warrants further investigation. In addition, GIV functions downstream of innate immune receptors, including TLR4^50^ and NOD2^57^, where it has established roles in regulating inflammatory signaling^90,112–118^; the extent to which the GIV checkpoint coordinately integrates lipid handling and inflammatory programs within atherosclerotic lesions remains unresolved. Limited bioavailability of IGGi-11me, presumably due to high first-pass metabolism^67^ restricted its evaluation in mice; medicinal chemistry efforts are necessary before efficacy and safety can be rigorously assessed. Structural definition of the GIV•ABCA1 interface (e.g., cryo-EM) will also be important for structure-guided therapeutic design. Finally, the mechanism linking GIV loss to enhanced lipolysis remains unclear, but may involve elevated autophagic flux, a known feature of GIV-deficient cells^119^, or high cAMP levels, which favor lipolysis^94^.

## Conclusions

By identifying GIV as a master brake on cholesterol transport, this work reframes macrophage lipid biology as a tractable therapeutic frontier. In contrast to statins and β-blockers, which stabilize but do not “defat,” GIV inhibition converges on the root cause, i.e., lipid persistence. Targeting this noncanonical G-protein checkpoint offers a unified molecular and pharmacologic strategy to reprogram macrophages, restore RCT, and accelerate the long-sought regression of atherosclerosis.

## Resource Availability Lead Contact

Further information and requests for resources and reagents should be directed to and will be fulfilled by the lead contact, Pradipta Ghosh.

## Materials Availability

All materials are available from the lead contact with a completed Materials Transfer Agreement and patented technology agreement following the guidelines of the University of California, San Diego.

## Data and Code Availability

All materials are available from the lead contact with a completed Materials Transfer Agreement and patented technology agreement following the guidelines of the University of California, San Diego. All transcriptomic datasets generated in this work has been deposited at NCBI GEO [GSE309374 and GSE309324]. The data underlying all the Figures and tables are available in the article and its online supplemental materials. The Boolean analysis framework is available at https://github.com/sinha7290/Atherosclerosis. Any additional information required for reanalysing the data reported in this work paper is available from the Lead Contact upon request.

## Acknowledgments

This work was supported by the National Institutes of Health (NIH) (grant# R01-AI141630 and R01-AI55696 to P.G.). G.D.K. received support from the American Association of Immunologists Intersect Fellowship Program for Computational Scientists and Immunologists. Both G.D.K and M.S.A were also supported in part by the American Heart Association (24CDA1268506 and 25PRE1357971, respectively). S.S. was supported by the AAI Intersect Fellowship Program. M.M. was supported by a UC San Diego Agilent Center of Excellence Postdoctoral Fellowship. Data in this manuscript were generated at the UC San Diego Institute of Genomic Medicine using an Illumina NovaSeq 6000, funded by NIH SIG grant S10-OD026929. The authors acknowledge instrumentation resources at the UC San Diego Agilent Center of Excellence in Cellular Intelligence. The authors acknowledge that lipid analysis was performed at the UCSD Lipidomics Core.

The content is solely the authors’ responsibility and does not necessarily represent the official views of the Helmsley Charitable Trust, AHA or the NIH.

## Author contributions

G.D.K. and P. G. conceptualized the project. C.R.E. generated the *Apoe*^-/-^ *Ccdc88a*^fl/fl^ *LysM*^Cre/-^ and *Apoe*^-/-^*Ccdc88a*^fl/fl^ mouse models, designed, cloned, and performed site-directed mutagenesis of all constructs used in this work. S.S., with supervision from P.G., performed all transcriptomic analyses. G.D.K., M.S.A., C.R.E., M.E., C.B., M.N., E.T., R.M., V.C., Y.S.M., M.M., and B.K. contributed to data curation and formal analysis. G.D.K., V.C., R.M., and B.K. conducted all animal studies. A.A., and Q.O., conducted mice fecal sterol analysis. M.S.A. conducted all biochemical studies. C.R.E., M.M., and S.A., M.S.A. assisted with animal studies. M.E., C.B., E.M., and R.M. assisted G.D.K. with in vitro assays and analysis. J.Y., supervised the synthesis of chemical matter. G.D.K. and P.G. prepared the Figures for data visualization. PG wrote the original draft. G.D.K., M.S.A., S.S., C.R.E., and P.G. reviewed and edited the manuscript. P.G. supervised and acquired funding to support the study. All co-authors approved the final version of the manuscript.

## GENERATIVE AI USE DISCLOSURE

Generative AI has not been used to write the manuscript or generate data figures de novo. AI-related tools were used to create schematics or to improve the readability and language of the text after it had been drafted.

### STAR ★ Methods

#### Key resources table

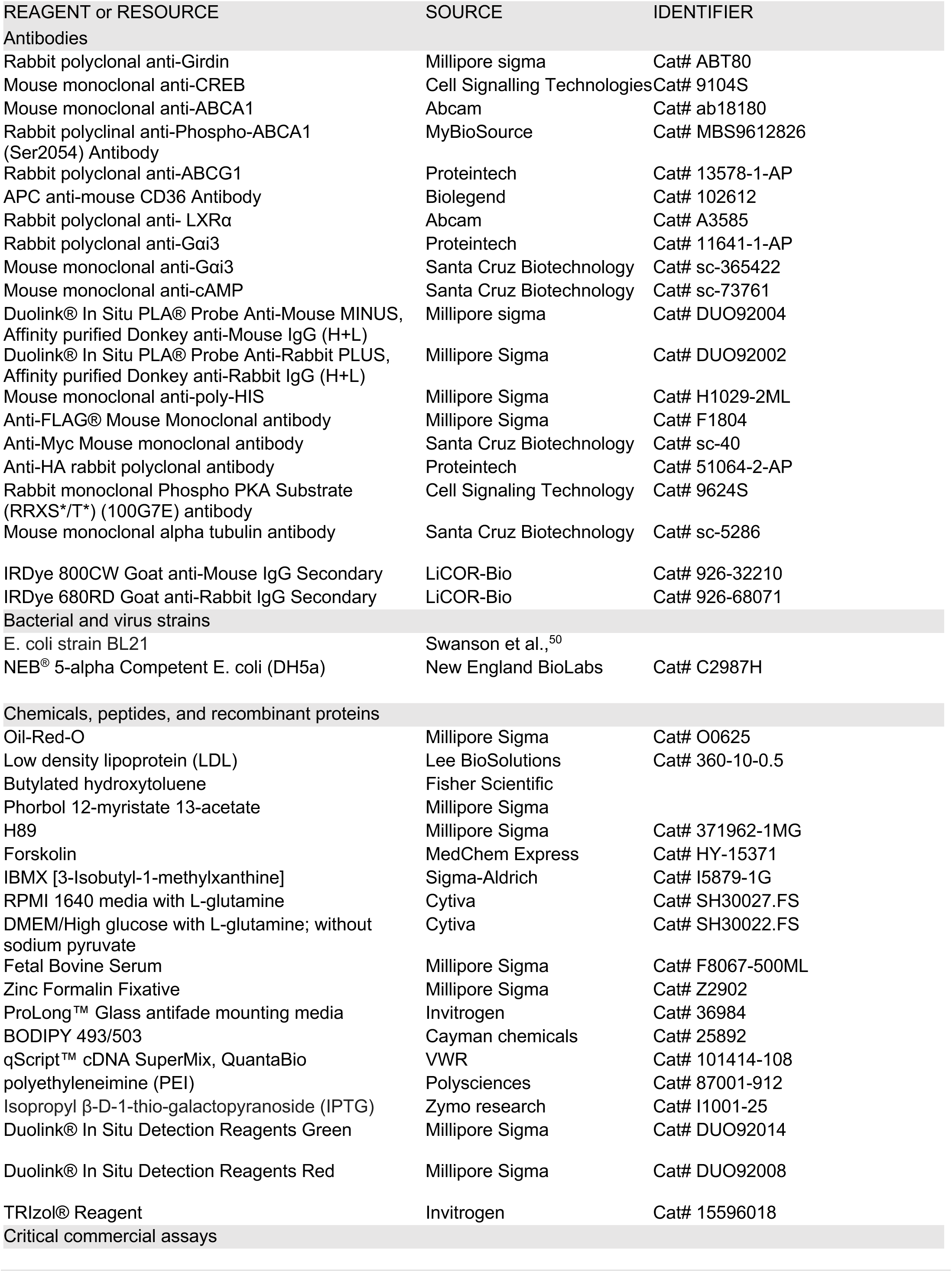
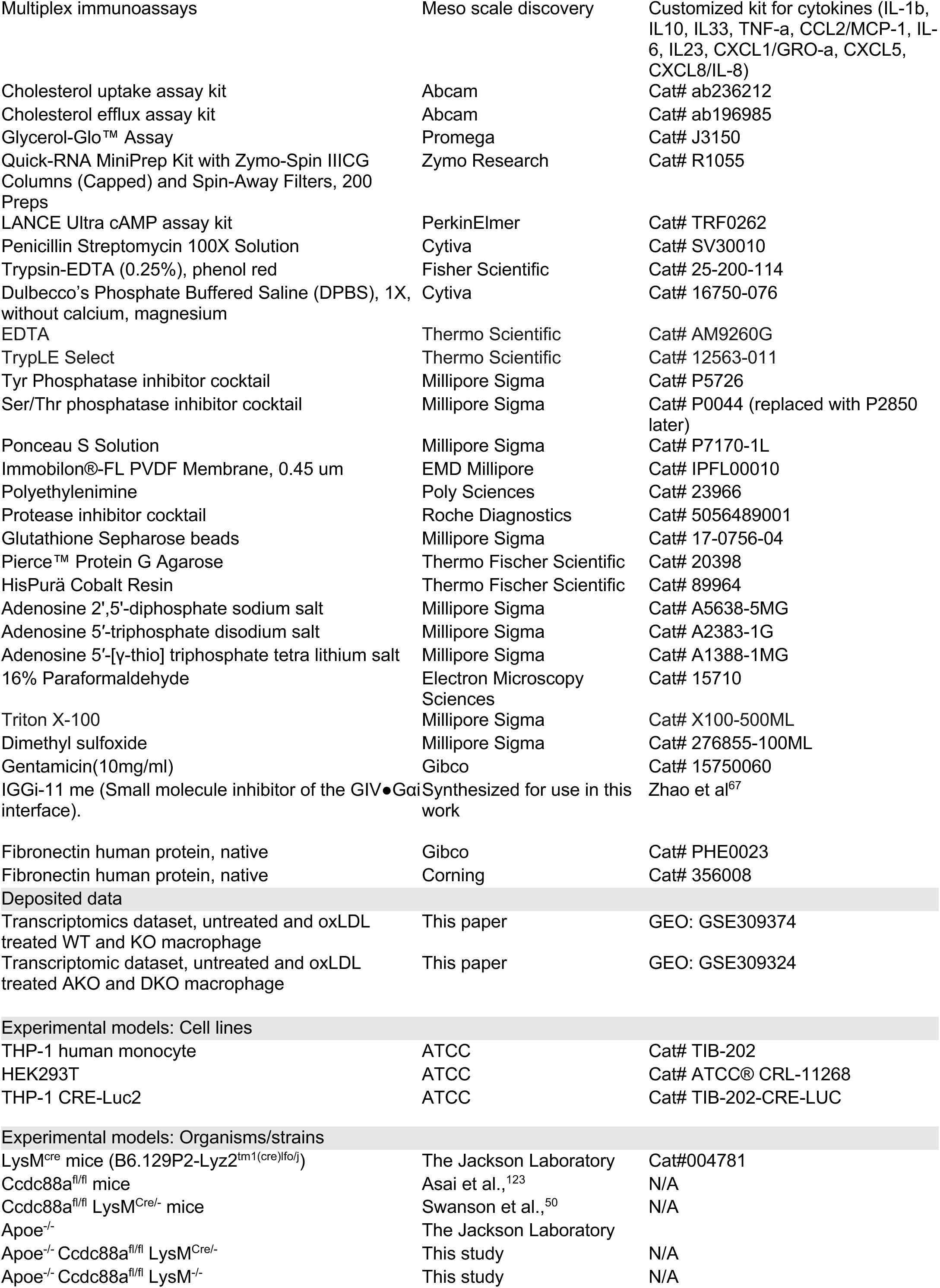
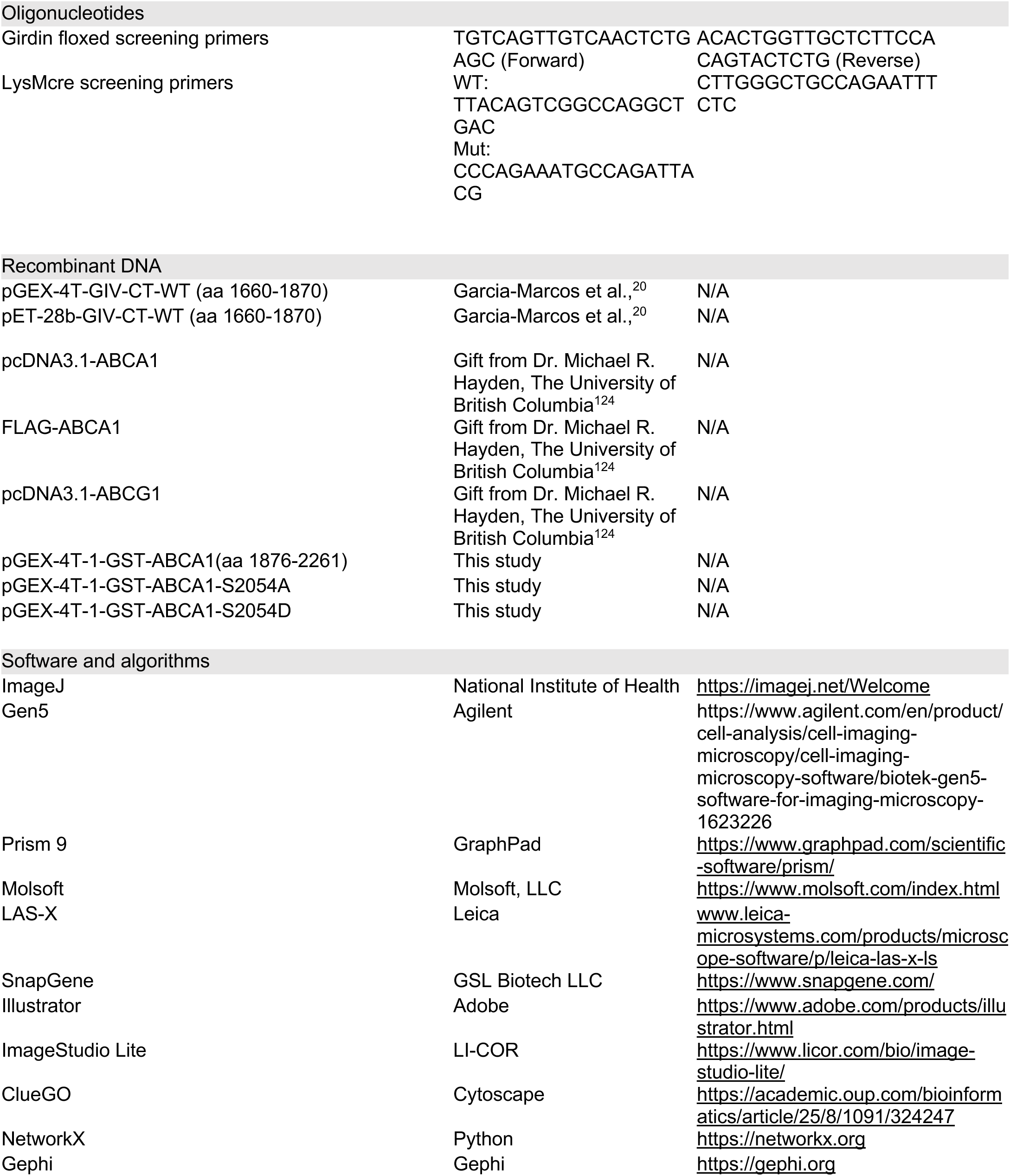

### Experimental models and Study Participant Details

#### Cell

Cell lines used in this study were obtained from either American Type Culture Collection (ATCC) or InvivoGen. All lines were routinely tested for mycoplasma contamination using commercially available PCR-based kits. Only validated, mycoplasma-free cells were expanded and used for experiments.

#### THP-1

This human monocytic cell line was obtained from the ATCC and maintained in Roswell Park Memorial Institute 1640 (RPMI-1640) medium (Gibco-BRL). The medium was supplemented with heat-inactivated 10% fetal calf serum (Hyclone), 1% L-glutamine, and 1% penicillin/streptomycin (Gibco-BRL). The cells were incubated at 37°C in a 5% CO₂ incubator. Cultures were subcultured by centrifugation and resuspension at 2–4 × 10⁵ viable cells/ml, and maintained below 1 × 10⁶ cells/ml.

#### THP1-CRE-Luc

This CREB reporter derivative of the THP-1 cell line was obtained from InvivoGen and maintained in RPMI-1640 medium supplemented with heat-inactivated 10-20% fetal calf serum, 1% L-glutamine, and 1% penicillin/streptomycin, as per the manufacturer’s direction. The cells were incubated at 37°C in a 5% CO₂ incubator. Cultures were subcultured by centrifugation and resuspension at 2–4 × 10⁵ viable cells/ml, and maintained below 1 × 10⁶ cells/ml.

#### HEK293T

This human embryonic kidney cell line was obtained from the ATCC and maintained in Dulbecco’s Modified Eagle Medium (DMEM; Gibco-BRL) supplemented with 10% fetal calf serum, 1% L-glutamine, and 1% penicillin/streptomycin. The cells were incubated at 37°C in a humidified atmosphere with 5% CO₂ and were routinely passaged at a dilution of 1:5 to 1:10 using 0.05% trypsin-EDTA when they reached approximately 80–90% confluency.

#### Thioglycolate-elicited murine peritoneal macrophages

Murine peritoneal macrophages were collected from peritoneal lavage of 8- to 12-wk-old C57BL/6 mice with ice-cold RPMI (10 ml per mouse) 4 days after intraperitoneal injection of 3 ml of aged, sterile 3% thioglycolate broth (Becton Dickinson Difco™ Detroit, MI, USA) and cultured as described previously^125^. Peritoneal macrophages were passed through 70 µm filter to remove possible tissue debris contamination during harvesting. They were counted, centrifuged, and resuspended in RPMI-1640 containing 10% FBS and 1% penicillin/streptomycin and plated with the required cell density. The media was changed after 4 h to remove non-adherent cells. Macrophages were allowed to adjust to overnight culture before the addition of oxLDL (100 μg/ml) as indicated in the figure legends.

### Animal Model

*Ccdc88a^fl/fl^* mice were a gift from Dr. Masahide Takahashi (Nagoya University, Japan) and was developed as described^123^. *LysM*^Cre/Cre^ mice (B6.129P2-Lyz2tm1(cre)lfo/j) were purchased from the Jackson Laboratory. *Ccdc88a*^fl/fl^ x *LysM*^Cre/-^ mice were generated previously by us as described^50^ and were maintained as homozygous floxed and heterozygous LysMcre. The *Apoe^-/-^*mice were purchased from the Jackson Laboratory. They were crossed with *Ccdc88a^fl/fl^*mice to generate *Apoe^-/-^ Ccdc88a*^fl/fl^ mice. The *Apoe^-/-^ Ccdc88a*^fl/fl^ mice were crossed to *LysM^cre/^*^cre^ *Ccdc88a^fl/fl^* mice to generate the *LysM^cre/^*^-^ *Apoe^-/-^ Ccdc88a*^fl/fl^ mice. Primers required for genotyping are mentioned in primer (*Key Resource Table*). All mice studies were approved by the University of California (UC), San Diego Institutional Animal Care and Use Committee (IACUC). Male mice (8-12 weeks) were used for all experiments and maintained in an institutional animal care at the UC San Diego animal facility on a 12-h/12-h light/dark cycle (humidity 30–70% and room temperature controlled between 68–75 °F) with free access to normal chow food and water. To promote atherosclerosis, animals were fed with *ad libitum* western diet (consisting of 60% fat, VWR, West Chester, PA, USA) for 3 months.

### Ethics Statement

All animal studies were conducted in strict accordance with the NIH *Guide for the Care and Use of Laboratory Animals* and were approved by the Institutional Animal Care (#S17223; PI Ghosh). All human studies were conducted at UC San Diego, with informed consent under an IRB-approved protocol (#190105; PI Ghosh).

## Method Details

### Computational Methods

#### Curation of transcriptomic datasets

##### Computational modeling of LAM heterogeneity

To resolve the transcriptional diversity of LAM, we applied the previously established Signatures of Macrophage Reactivity and Tolerance (SMaRT) model^44^, which captures macrophage continuum states across >12,500 datasets. The model was trained using bulk RNA-seq data from stable vs. complicated human carotid plaques (GSE120521; *n* = 8). Classification performance was evaluated by ROC analysis (AUC = 1.0, *p* < 0.05). To evaluate classification strength and prediction accuracy, Receiver Operating Characteristic (ROC) curves were generated for each gene. These curves assess the performance of a binary classifier, e.g., high vs. low *StepMiner* normalized gene expression levels, across varying discrimination thresholds. ROC curves plot the True Positive Rate against the False Positive Rate at multiple threshold levels. The Area Under the Curve (AUC) quantifies the classifier’s ability to correctly distinguish between the various groups of samples. ROC-AUC values were computed using the Python Scikit-learn package. Genes contributing significantly to model accuracy were designated LAM-associated SMaRT genes (n = 24).

##### StepMiner Algorithm

*StepMiner* is an algorithm designed to detect stepwise transitions in time-series gene expression data^126^. It fits step functions to expression profiles by identifying the sharpest changes in signal, corresponding to gene expression switching events. The algorithm evaluates all possible step positions and calculates the average expression on either side of each step to define constant segments. An adaptive regression approach is then used to select the step position that minimizes the sum of squared errors between the observed and fitted data. The selected step is used as the *StepMiner* threshold*. This* threshold is used to convert gene expression values into Boolean values (0 or 1). A noise margin of 2-fold change is applied around the threshold to determine intermediate values, and these values are ignored during Boolean analysis. Finally, a regression test statistic is computed to assess the significance of the identified step transition as follows:

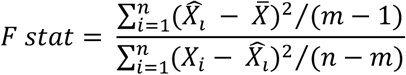

Where *X_i_* for *i* = 1 to *n* are the values, 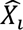 for *i* = 1 to *n* are fitted values. m is the degrees of freedom used for the adaptive regression analysis. 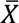 is the average of all the values: 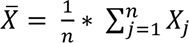. For a step position at k, the fitted values 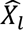 are computed by using 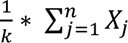 for *i* = 1 to *k* and 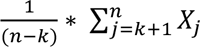 for *i* = *k* + 1 to *n*.

##### Composite Gene Signature Analysis Using Boolean Network Explorer

Boolean network explorer (BoNE)^127^ provides an integrated platform for the construction, visualization and querying of a gene expression signature underlying a disease or a biological process in three steps: **(i)** *<u>Binarization</u>*: The expression levels of all genes in these datasets are converted to binary values (high or low) using the *StepMiner* algorithm^126^. **(ii)** *<u>Normalization</u>*: Gene expression values are normalized using a modified Z-score approach centered on the *StepMiner* threshold [formula = (expr - SThr)/(3*stddev)]. **(iii)** *<u>Scoring</u>*: Normalized expression values for each gene in a given signature, are summed to generate a composite score, for the gene signature, representing the overall activity of the biological pathway in a sample.

As a modified Z-score, the composite score of a gene signature ranges from negative to positive values, reflecting the dynamic range of each gene in the signature. These composite scores enable the comparison of biological states between control and query groups within a dataset. However, comparisons between different gene signatures or across datasets are not valid, due to the lack of cross-signature normalization and dataset-specific normalization processes.

Samples were ranked by composite signature scores, and statistical differences between groups were assessed using Welch’s t-test (unpaired, unequal variances and sample sizes). Visualizations, including violin, swarm, and bubble plots, were generated using the Seaborn Python package (version 0.10.1).

##### RNA Sequencing and Data Processing

RNA sequencing libraries were generated using the Illumina TruSeq Stranded Total RNA Library Prep Gold with TruSeq Unique Dual Indexes (Illumina, San Diego, CA). Samples were processed following the manufacturer’s instructions, except for modifying the RNA shear time to five minutes. The resulting libraries were multiplexed and sequenced with 100 base pairs (bp) Paired-End (PE100) to a depth of approximately 40 million reads per sample on an Illumina NovaSeq 6000. Samples were demultiplexed using bcl2fastq v2.20 Conversion Software (Illumina, San Diego, CA). Raw FASTQ files were trimmed, filtered, and mapped to the human genome for downstream quantification analyses. Low-quality sequences were trimmed or removed with Trimmomatic. STAR (version 2.6.0a) was used to align reads on the reference genome (*Mus musculus* genome build GRCm38.94). The resulting transcriptome-aligned sequences were used for expression quantification by using RSEM (version 1.3.3) with “–forward-prob 0” option. TPM scores for each sample were used through the RSEM tables (RSEM gene. Results tables). We used log2(TPM) if TPM >1, else TPM - 1 values for each sample as the final gene expression value for downstream expression analyses.

##### Differential Expression Analysis

Differential gene expression analysis was performed using raw count data processed through DESeq2^128^. Genes with an absolute log2 fold change ≥ 0.5 and an adjusted p-value ≤ 0.05 were considered differentially expressed genes (DEGs). Pathway enrichment analysis of DEG lists was conducted using the ShinyGO ^129^ database and its integrated algorithms.

##### Expression Heatmap

Z-normalized expression values for selected genes were visualized as a heatmap across all samples within the dataset using the Seaborn package (version 0.10.1).

##### Odds calculation on clinical parameters and gene signatures

Clinical, treatment, and transcriptomics data were obtained from Doeing et. al.,^130^. Clinical and treatment parameters were compared between patients with good (stable) and bad (unstable/ruptured plaques) outcomes. For categorical variables, proportions were converted to counts based on group sizes (n=15 per group) and analyzed using 2×2 contingency tables. Odds ratios (ORs) with 95% confidence intervals (CIs) were calculated to estimate the strength of association between each parameter and outcome, and Fisher’s exact test was applied to assess statistical significance given the small sample size. Continuous variables (e.g., age) were compared descriptively using mean ± standard deviation and were not included in OR calculations. For stratification of patients using gene signatures, patients were divided into High vs Low signature groups based on median value of composite score, and the association with outcome was assessed in the same way using odds ratios, 95% CIs, and Fisher’s exact test.

#### Experimental

##### Oil-Red-O (ORO) staining of aorta and quantification of plaque burden

ORO staining was done as previously described^131^. Briefly, murine aortas were isolated, perfused, fixed in 4% paraformaldehyde (PFA) and stained with freshly prepared and filtered 0.3% ORO solution for 60 min at room temperature, with continuous rotation. Stained aortas were imaged using a Leica DM1000 light microscope (Leica, Vista, CA, USA) or Biotek Cytation 10 (Agilent Technologies - La Jolla, CA, USA), and analysis was conducted using ImageJ (NIH, Bethesda, MD, USA) or Gen5 software (Agilent Technologies - La Jolla, CA, USA).

##### Histology analysis

Randomly selected 7-8 samples of dissected liver and adipose tissues were fixed in 4% paraformaldehyde (PFA) and embedded in paraffin blocks. Sections of 5-7 μm thickness were prepared and stained with hematoxylin and eosin (H&E). Lipid accumulation was quantified from the H&E-stained images using Gen5 software and normalized to the respective H&E-stained areas.

##### Total Sterol/Oxysterol analysis

Total sterols and oxysterols in mouse feces were quantified using established protocols^132,133^ with minor modifications. Briefly, mouse fecal pellets weighing approximately 100mg were homogenized in 1 ml of 10% methanol in water. An internal standard mix of 25-Hydroxycholesterol-d6, Desmosterol-d6, and Campesterol-d6 (Avanti Polar Lipids) was added to 250 μl of the homogenate. Samples were saponified for 1.5 h at 37 °C with a final concentration of 0.5 N KOH. Samples were extracted with 500 μl of butanol/methanol (3:1, v/v), heptane/ethyl acetate (3:1, v/v), and 1% acetic acid in water (BUME). Extracts were brought to dryness and taken up in 90% methanol in water and run on a Waters Acquity UPLC interfaced with an AB Sciex 6500 QTrap mass spectrometer equipped with an APCI probe. Source settings were: Curtain Gas=20, Collision Gas=Medium, Ion Spray Voltage=5500, Temperature=400, GS1=25, NC=1. A Phenomenex Kinetex C18 1.7 μm 2.1 mm x 150 mm column was used for chromatographic separation. A 30 min step gradient was employed using 70/30 acetonitrile/water with 5 mM ammonium acetate as Buffer A and 50/50 acetonitrile/water with 5 mM ammonium acetate as Buffer B with a flow of 0.5 ml/min. Sterol species were identified by mass spectrometry using 30 MRMs (Multiple Reaction Monitoring) in positive mode (see Data S4). Standard curves were obtained in parallel using identical conditions. Data analysis was performed with Analyst and Multiquant software packages.

##### Oxidized low density lipoprotein (oxLDL) preparation

The oxLDL was prepared as previously described^134^. Briefly, oxidized LDL (oxLDL) was generated by incubating LDL (2 mg/ml in PBS; 360–10, Lee BioSolution, Maryland Heights, MO, USA) with 10 µM CuSO₄ at 37 °C for 18 h, and oxidation was halted by the addition of 20 µM butylated hydroxytoluene and 300 µM EDTA.

##### LAM formation and quantification

LAM formation was assessed as previously described^125^ with minor modifications. Peritoneal macrophages were incubated with freshly prepared 50/100 μg/ml oxLDL for 0-24 as indicated in the figure legends. Cells were then fixed with 4% PFA and stained with freshly filtered 0.3% Oil-Red-O (ORO) working solution for 60 min at room temperature. For THP1 cells, upon oxLDL treatment cells, were fixed and stained with BODIPY (2μM)/DAPI (1:500 dilution). Coverslips were mounted using ProLong™ Glass antifade (Invitrogen Carlsbad, CA, USA) and used for imaging. Cells were imaged using Stellaris confocal microscope (Leica, Vista, CA, USA) or Biotek Cytation 10 and analysis was conducted using ImageJ or Gen5 software.

##### Cholesterol uptake assay

A Cholesterol Uptake Assay Kit (Abcam, Cambridge, UK), following the manufacturer’s guidelines. Briefly, peritoneal macrophages (40,000 cells/well) were incubated with fluorescently labeled NBD-Cholesterol for 24 h in serum-free RPMI media. After incubation, the macrophages were treated with assay buffer for an additional 24 h. The supernatant was then collected, and fluorescence was measured using the FITC filter on the Spark® 20M multimode reader (Tecan, Morrisville, NC, USA).

##### Lipolysis Assay

A Glycerol-Glo™ Assay kit (Promega, J3150) was used, according to the manufacturer’s instructions with minor modifications. Briefly, cell culture media supernatant collected after oxLDL treatment was used to assess glycerol release as a readout of lipolytic activity. Supernatants (25 μl) were mixed with an equal volume of Glycerol Detection Reagent, incubated for 30 min at room temperature, and luminescence was measured using a Spark® 20M multimode reader. Results were expressed as relative glycerol release or fold-change over control.

##### Cholesterol efflux assay

A Cholesterol Efflux Assay kit (Abcam, Cambridge, UK) was used, as per the manufacturer’s guidelines. Briefly, peritoneal macrophages (40,000 cells/well) were labelled with fluorescent cholesterol (Ex/Em 485/523 nm) and treated with an equilibration reagent mix overnight to prevent fluorophore-labelled cholesterol breakdown. The following day, after washing, cholesterol efflux was initiated by adding samples containing cholesterol acceptor apolipoproteins. After a 4 h incubation, the supernatant (medium with cholesterol acceptors) was transferred to a white 96-well plate, and fluorescence (Ex/Em = 485/523 nm) was measured in endpoint mode. To quantify the cholesterol efflux, the adherent macrophage cell monolayer was solubilized with lysis buffer, and the lysate was transferred to another white 96-well plate for fluorescence measurement. The efflux from the labelled macrophages to a specific cholesterol acceptor was calculated by dividing the fluorescence intensity (RFU) of the supernatant by the sum of the fluorescence intensities of both the supernatant and the cell lysate from the same treatment.

##### Cytokine profile

Supernatants of cells treated with or without oxLDL were analyzed for the presence of various cytokines using customized Meso Scale Discovery (MSD) V-PLEX cytokine panels according to the manufacturer’s instructions. Data were collected with a MESO QuickPlex SQ 120 instrument (see *Key Resources Table*) and analyzed using MSD Discovery Workbench 4.0 software (see *Key Resources Table*).

##### General notes for the synthesis and characterization of IGGi-11me

IGGi-11me was synthesized and purified following the procedures described by Mahitha et al. Briefly, the multistep synthesis involved preparation of sodium *9H-fluorene-2,7-disulfonate*, conversion to the disulfonyl dichloride, synthesis of *4-(methylamino) methyl butanoate*, and final coupling to obtain *dimethyl 4,4’-((9H-fluorene-2,7-disulfonyl)bis (methylazanediyl)) dibutyrate* (IGGi-11me). The product was purified by column chromatography and preparative HPLC.

##### RNA isolation

RNA was isolated from foam cells using Direct-zol RNA Miniprep Kits, following the manufacturer’s protocol. Briefly, foam cells were thoroughly homogenized in Tri-reagent and processed through Zymo-Spin™ Column, followed by DNase-I digestion to purify total RNA. RNA was eluted in 50 μl free water. The quality of RNA was assessed using a Nanodrop instrument.

##### Flow cytometry

For macrophage cell surface expression analysis, cells were detached using trypsinization, washed with phosphate-buffered saline (PBS), and blocked for 10 min at room temperature in PBS containing 2 mg/ml bovine serum albumin (BSA). Cells were then incubated for 30 min at room temperature with fluorescently conjugated or unconjugated primary antibodies targeting ABCA1 (Abcam, Cambridge, UK), ABCG1 (Proteintech, Rosemont, IL, USA), and CD36 (BioLegend, La Jolla, CA, USA). After washing, if primary antibodies were unconjugated, cells were incubated with a fluorescent-conjugated secondary antibody (Invitrogen Carlsbad CA, USA, dilution 1:500 in BSA/PBS) for an additional 30 min at room temperature. Unstained samples were used as controls. For detecting total expression of the target proteins, cells were fixed and permeabilized before staining, following the same antibody incubation steps. Flow cytometry was performed using the Agilent Penteon or Opteon, a high-parameter spectral flow cytometer (Agilent Technologies - La Jolla, CA, USA). Data acquisition and analysis were conducted using the Agilent NovoExpress software (Agilent Technologies - La Jolla, CA, USA), ensuring precise quantification of surface and intracellular protein expression.

##### Cyclic AMP quantification

Cyclic AMP (cAMP) levels were determined using the LANCE Ultra cAMP Assay kit (PerkinElmer, Waltham, MA, USA). Peritoneal macrophages (5000 cells/well) were seeded in a 96-well low-volume plate and were either kept untreated or treated with 0.1 μg/ml oxLDL for 30 min. Following incubation, Eu-cAMP and U*Light* anti-cAMP antibody were added in each well and incubated at RT. After 1 h, FRET emission was recorded using Spark® 20M multimode reader.

##### Quantitative immunoblotting

Whole cell lysates were prepared by lysing cells in HEPES lysis buffer [20 mM HEPES (pH 7.2), 5 mM magnesium acetate, 125 mM potassium acetate, 0.4% Triton X-100, 1 mM DTT] supplemented with 500 µM sodium orthovanadate, phosphatase inhibitor cocktail (Sigma-Aldrich), and protease inhibitor cocktail (Roche, California, USA). Lysates were passed through a 28 G needle on ice and centrifuged at 10,000 × g for 10 min at 4°C.

For immunoblotting, the boiled protein samples were separated by 8- 10% SDS-PAGE and transferred to polyvinylidene fluoride (PVDF) membranes. Membranes were blocked with 5% non-fat milk in Tris-buffered saline with 0.1% Tween® 20 detergent (1X TBST). The membrane was stained with Ponceau S to visualize bait proteins (GST), washed, blocked (5% milk), and incubated with primary antibody solutions overnight at 4°C (unless otherwise specified, a 1:500 dilution for primary antibody was used). Phospho-PKA activity was assessed by immunoblotting with a rabbit monoclonal anti-phospho-PKA substrate (RRXS*/T*) antibody (Cell Signaling Technology, Danvers, MA, USA; 1:1000 dilution). Mouse monoclonal α-Tubulin (Santa Cruz Biotechnology, Dallas, TX, USA; 1:1000 dilution) served as a loading control. Washed blots were then incubated with infrared secondary antibodies (see *Key Resource Table*) for 1 h at room temperature.

All immunoblots were imaged and quantified using a LI-COR Odyssey imaging system (LI-COR Bioscience, Lincoln, NE, USA.) with dual-color infrared detection using its densitometry feature and analyzed using the Image Studio Lite software as per the manufacturer’s instructions.

Bound proteins were normalized to soluble proteins and % bound was expressed as fold change compared to internal controls on the same immunoblot to avoid confounding variables such as antibody potency, transfer efficiency, etc.

##### Assessment of total and phosphorylated ABCA1

THP1-derived serum synchronized macrophages were stimulated with 100 µg/mL oxLDL for 60 min were lysed in RIPA buffer [20 mM HEPES (pH 7.2), 1 mM EDTA, 180 mM NaCl, 0.5% sodium deoxycholate, 1% Triton X-100, 0.1% SDS] supplemented with sodium orthovanadate, phosphatase inhibitors (Sigma), and protease inhibitors (Roche). Lysates were passed through a 28-gauge needle on ice, cleared by centrifugation (10,000 × g, 10 minutes, 4 °C), and warmed at 42 °C for 10 min prior to SDS-PAGE. Phosphorylated and total ABCA1 were detected using rabbit polyclonal anti-pS^2054^–ABCA1 (MyBioSource; San Diego, CA; USA 1:1000 dilution) and mouse monoclonal total ABCA1 (Abcam; Cambridge; UK; 1:1000 dilution) antibodies. Subsequent immunoblotting steps, including membrane blocking, antibody incubation, infrared secondary detection, imaging, and densitometric quantification, were performed as described in the Quantitative immunoblotting section.

##### Confocal Immunofluorescence

Cells were fixed with 4% paraformaldehyde in PBS for 30 min at room temperature, followed by treatment with 0.1 M glycine for 10 min to quench autofluorescence. Blocking and permeabilization were performed using PBS containing 1% BSA and 0.1% Triton X-100 for 20 min. Primary antibodies (CREB (1:250), Cell Signaling Technologies, Danvers, MA, USA) were incubated for 1 h at room temperature or overnight in blocking buffer, followed by Alexa Fluor-conjugated secondary antibodies (1:500) for 1 h. For neutral lipid visualization, 1 μM of BODIPY stain (Cayman chemicals, Ann Arbor, MI, USA) was added along with secondary antibodies. After three PBS washes, coverslips were mounted using ProLong Glass antifade reagent for imaging. Cells were imaged using Stellaris confocal microscope or Biotek Cytation 10 and analysis was conducted using ImageJ or Gen5 software.

##### Plasmid constructs

Cloning of GIV-CT (1660-1870) into pET28b (His-GIV CT) were previously described^50^. The nucleotide region of the human ABCA1 gene encoding amino acids 1876–2261 (NM_005502.4/NP_005493.2), generously provided by Michael R. Hayden, was PCR-amplified from the pcDNA3.1-ABCA1 plasmid using Q5 High-Fidelity DNA Polymerase (New England Biolabs, Ipswich, MA). The PCR product was ligated into the SmaI/XhoI restriction sites of the pGEX-4T-1 vector to generate the GST-ABCA1(1876–2261) construct. ABCA1 mutant plasmids (GST-ABCA1-S2054A, and GST-ABCA1-S2054D) were generated by site directed mutagenesis on GST-ABCA1(1876-2261) using the Q5Ò Site-Directed Mutagenesis Kit (New England Biolabs, Ipswich, MA) according to manufacturer’s protocol. Plasmid constructs and their mutants were verified by DNA sequencing (Genewiz, La Jolla, CA), and protein expression was confirmed by western blot analysis.

##### Transfection, lysis, and co-immunoprecipitation

HEK293T were cultured in DMEM media containing 10% FBS and antibiotics, following ATCC guidelines. Cells were transfected with DNA plasmids using polyethyleneimine (PEI) following the manufacturer’s protocols^135^. HEK293T cells were expressed with either N-terminal FLAG-tagged or untagged full-length ABCA1^9^. 48 h post-transfection, cells were lysed in buffer containing 20 mM HEPES, pH 7.2, 5 mM Mg-acetate, 125 mM K-acetate, 0.4% Triton X-100, 1 mM DTT, 0.5 mM sodium orthovanadate, and phosphatase (for both Tyr and Ser/Thr) and protease inhibitor cocktails.

For immunoprecipitation, equal aliquots of clarified cell lysates were incubated with 2 μg of anti-ABCA1 antibody (Anti-ABCA1 antibody, ab18180, Abcam, Cambridge, UK) for 3 h at 4°C, followed by the addition of protein G Sepharose beads (GE Healthcare; 40 µl 50% v:v slurry) and incubation for 1 h at 4°C. Beads were washed four times (1 mL volume each wash) in PBS-T buffer [4.3 mM Na2HPO4, 1.4 mM KH2PO4, pH 7.4, 137 mM NaCl, 2.7 mM KCl, 0.1% (v:v) Tween 20, 10 mM MgCl2, 5 mM EDTA, 2 mM DTT, 0.5 mM sodium orthovanadate] and immune complexes were eluted by boiling in Laemmli’s sample buffer.

Elutes were resolved on SDS PAGE and analyzed by immunoblotting with mouse anti-ABCA1 (1:500 dilution), rabbit anti-LXRα (Abcam, ab3585, 1:1000 dilution) and rabbit anti-GIV antibody (ABT80, Millipore sigma, Burlington, MA, 1:500 dilution)

### Single and sequential DuoLink® Proximity Ligation Assay (PLA)

PMA-differentiated THP-1 wild-type (WT) cells were seeded on sterile glass coverslips at a density sufficient to reach ∼60–70% confluency and treated with 25 nM phorbol 12-myristate 13-acetate (PMA; Sigma-Aldrich, St. Louis, MO) for 16–18 h to induce differentiation. The next day, cells were either left unstimulated or stimulated with 50 µg/mL OxLDL for 24 h. Following stimulation, cells were washed twice with phosphate-buffered saline (PBS) and fixed with 4% paraformaldehyde (PFA, in PBS) for 20 min at room temperature (RT), quenched with 0.1 M glycine in PBS for 10 min, permeabilized with 0.2% Triton X-100 in PBS for 1 h at RT and blocked with 1% bovine serum albumin (BSA) and 0.1% Triton X-100 in PBS for 1 h, as described previously^56^.

Protein–protein interactions were assessed using the Duolink® In Situ Red and Green PLA kits (Sigma-Aldrich) following the manufacturer’s instructions. For single PLA assay, coverslips were incubated overnight at 4°C with primary antibodies against GIV (rabbit, Millipore; ABT80, 1:150 dilution) and ABCA1 (mouse, Abcam, ab18180,1:50 dilution), followed by incubation with species-specific PLA probes (PLUS and MINUS) for 1 h at 37°C in a humidified chamber. Ligation and rolling circle amplification were performed. Red PLA detection reagents (Ex/Em: 566–594 nm) visualized GIV●ABCA1 interactions, ABCA1●LXR and ABCA1●Gαi3 interactions. For GIV●Gαi3 detection, rabbit anti-GIV (1:150) and mouse anti-Gαi3 (1:50) ; were used, followed by Green PLA detection reagents (Ex/Em: ∼488 nm). Each individual PLA interaction was performed on separate cover slips—in parallel—from the same experiment but imaged, analyzed, and quantified independently.

For the dual PLA assay, GIV●ABCA1 interaction was visualized using the red PLA protocol. After PBS washes, coverslips were incubated overnight with mouse anti-Gαi3, and the green PLA protocol was repeated to detect GIV●Gαi3 interactions.

In all assays, nuclei were counterstained with 1 µg/mL DAPI to visualize nuclei and coverslips were mounted using ProLong™ Gold. Imaging was performed using Leica Stellaris confocal microscope or BioTek Cytation 10 (Agilent Technologies). PLA signal quantification (puncta/cell) was done in ImageJ using background subtraction, thresholding, and the “Analyze Particles” tool.

#### Protein expression and purification

His-tagged and GST-tagged recombinant proteins were expressed in *E. coli* strain BL21 and purified as previously described^50,136,137^. Bacterial cultures were induced with 1 mM isopropyl β-D-1-thio-galactopyranoside (IPTG) either overnight at 25°C. Bacterial culture (1l) were pelleted after induction and resuspended in 20 mL GST-lysis buffer (25 mM Tris·HCl, pH 7.5, 20 mM NaCl, 1 mM Ethylenediaminetetraacetic acid (EDTA), 20% [v/v] glycerol, 1% [v/v] Triton X-100, 2 × protease inhibitor mixture [Complete EDTA-free; Roche Diagnostics]) or in 20 mL His-lysis buffer (50 mM NaH_2_PO_4_ [pH 7.4], 300 mM NaCl, 10 mM imidazole, 1% [vol/vol] Triton X-100, 2 × protease inhibitor mixture [Complete EDTA-free; Roche Diagnostics]) for GST or His-fused proteins, respectively. Bacterial lysates were sonicated (three cycles, with pulses lasting 30 s/cycle, and with 2 min interval between cycles to prevent heating), centrifuged at 12,000×*g* for 20 min at 4°C. Supernatant (solubilized proteins) were affinity purified on glutathione-Sepharose 4B beads (GE Healthcare) or HisPur Cobalt Resin (Pierce), dialyzed overnight against PBS, and stored at −80°C.

#### GST pulldown assays

Recombinant GST-tagged proteins, including GST alone (control), GST-GIV-CT, GST-ABCA1 WT (aa 1876-2261) and its mutants, were expressed in *E. coli* strain BL21 (DE3) and purified as previously described^50^. Bacterial cultures were induced with 1 mM IPTG and incubated overnight at 25 °C. Cells were pelleted and resuspended in GST lysis buffer [25 mM Tris-HCl (pH 7.5), 20 mM NaCl, 1 mM EDTA, 20% (v/v) glycerol, 1% (v/v) Triton X-100, and 2× protease inhibitor cocktail], then briefly sonicated (30-second pulses on/off for 5 min). Lysates were clarified by centrifugation at 13,000 rpm for 45 min. Protein concentrations were determined using BSA standards, and aliquots were stored at –80 °C.

Equimolar concentrations of GST-tagged proteins were immobilized onto glutathione-Sepharose beads by incubation in binding buffer [50 mM Tris-HCl (pH 7.4), 100 mM NaCl, 0.4% (v/v) Nonidet P-40, 10 mM MgCl₂, 5 mM EDTA, 2 mM DTT, and protease inhibitors] for 1 h at 4 °C or overnight with gentle tumbling. After washing, the beads were incubated with equimolar amounts of purified His-tagged GIV-CT for 4 h at 4 °C. Ten percent of the His-tagged GIV CT input was retained as control.

Following incubation, beads were washed four times with phosphate wash buffer [4.3 mM Na₂HPO₄, 1.4 mM KH₂PO₄ (pH 7.4), 137 mM NaCl, 2.7 mM KCl, 0.1% (v/v) Tween-20, 10 mM MgCl₂, 5 mM EDTA, 2 mM DTT, 0.5 mM sodium orthovanadate], then eluted twice in Laemmli buffer [5% SDS, 156 mM Tris base, 25% glycerol, 0.025% bromophenol blue, 25% β-mercaptoethanol] by boiling at 95 °C for 5 min. Eluates were pooled, resolved by 10% SDS-PAGE, and transferred to PVDF membranes.

Membranes were stained with Ponceau S to confirm transfer efficiency and equimolar loading of GST-tagged proteins, then blocked in PBS containing 5% non-fat milk. Immunoblotting was performed using mouse anti-His antibody (Millipore Sigma, Burlington, MA, 1:1000 dilution), anti-FLAG (Millipore Sigma, 1:500 dilution) anti-ABCA1 (1:500 dilution). Detection and quantification were carried out using a LI-COR Odyssey imaging system with dual-color infrared detection. Final figures were assembled using Adobe Photoshop and Illustrator (Adobe, San Jose, CA, USA).

#### Human plaque-in-a-dish models

Peripheral blood samples were collected into heparinized tubes, and peripheral blood mononuclear cells (PBMCs) were isolated by density gradient centrifugation using HISTOPAQUE (Sigma-Aldrich, St. Louis, MO, USA). The PBMC layer was harvested using a Pasteur pipette, washed once with PBS, and pelleted by centrifugation at 250 × *g* for 10 min at 4 °C. The freshly purified PBMCs were subsequently used for in vitro experiments.

For plaque-in-a-dish assays, 12-well culture plates were coated with 500 µl of human plasma fibronectin (5 µg/ml; Corning Inc., Corning, NY) for 1 h at room temperature. The justification for the use of fibronectin is detailed in *Results*. Most importantly, we chose a source of fibronectin that would have minimal amounts of the alternatively spliced exon encoding type III repeat extra domain A (EDA), which is produced in response to tissue injury. This ensures that the confounding effects of EDA’s ability to stimulate TLR4^138^ and stabilize plaque^138^ are minimized. Plates were then washed once with sterile PBS and dried for 1 h at 37 °C. Freshly isolated PBMCs were seeded at a density of 1 × 10⁶ cells/1 ml per well onto fibronectin-coated plates and incubated for 3 h at 37 °C in a CO₂ incubator to allow monocyte adhesion. Non-adherent cells were removed after 4 h by media exchange, and adherent monocytes were cultured for ∼4–5 days to allow differentiation into adherent macrophages before being exposed to oxidized LDL (oxLDL, 100 µg/ml) for 24 h to create lipid-associated macrophages.

Cells were detached using TrypLE, centrifuged and washed with ice-cold sterile PBS. Neutral lipid staining was performed by incubating cells with BODIPY (1 µM) for 30 min at 4 °C, followed by two washes with FACS buffer and resuspension in 200 µl of FACS buffer. Samples were immediately analyzed on an Agilent Novocyte Penteon flow cytometer. The fraction of macrophages that transformed into lipid-associated macrophages ranged from ∼70-90%.

#### Software and statistical analysis

All images were processed using ImageJ software, Gen5, LAS X, or iStudio (LiCOR, Lincoln, NE, USA) software, and assembled into figure panels with Photoshop and Illustrator (Adobe Creative Cloud). All graphs were generated using GraphPad Prism v9.3.1. All experiments were repeated at least three times, and results were presented as average ± S.D. Statistical significance was assessed using either a t-test or one-way analysis of variance (ANOVA), including a Tukey’s test for multiple comparisons. Actual *p* values are displayed.

## Supplemental Information Index

### Supplemental Figures and Legends (Figure S1-S10)

Figure S1: Identification of *CCDC88A* as a putative target of lipid-associated macrophages, related to Figure 1.

Figure S2: Myeloid GIV loss reduces plaque burden without altering blood cholesterol and circulating myeloid populations, related to Figure 2.

Figure S3: GIV loss does not alter body, liver or adipose tissue mass in hyperlipidemic mouse models, related to Figure 2.

Figure S4: Fecal sterol profiling during diet-induced hyperlipidemia, related to Figure 2.

Figure S5: Survival of macrophages from diet-induced hyperlipidemia, related to Figure 3.

Figure S6: Validation of lipid-associated macrophage (LAM) and plaque stabilization gene signatures, related to Figure 4.

Figure S7: GIV scaffolds ABCA1 to Gαi, sequestering the transporter in endomembranes, related to Figure 5.

Figure S8: GIV’s C-terminus binds to ABCA1, related to Figure 6.

Figure S9: Pharmacogenomic evaluation of how the PKA-CREB pathway impacts LAM formation, related to Figure 7.

Figure S10: Impaired defatting in statin-exposed macrophages and counterbalancing pathways in atherosclerosis risk, related to Figure 8.

### Supplemental Tables (Table S1-S3)

Table S1: List of genes in the LAM gene signature, related to Figure 1B.

Table S2: The odds ratio of atherosclerosis based on clinical and molecular gene signatures, related to Figure 8I-K.

Table S3: The odds ratios for atherosclerosis based on risk factors, related to Figure 8 and S10.

### Supplemental Data (Data S1-S7)

Data S1: List of SMaRT genes, related to Figure 1.

Data S2: List of TREM2+ LAM gene signature, related to Figure 1. Data S3: Analysis of fecal sterols, related to Figure 2.

Data S4: Sterol and oxysterol standards, related to Figure 2.

Data S5: List of genes in a plaque-stabilization signature, related to Figure 4.

Data S6: List of genes in the athroprotective RCT gene signature, related to Figure 4. Data S7: Odds Ratio, related to Figure 8.

