## Supplementary information for "A Druggable G Protein Checkpoint in Cholesterol Efflux"

**Affiliations:**
[1]Department of Cellular and Molecular Medicine, University of California San Diego, CA, USA.
[2]Biomedical Sciences Graduate Program, School of Medicine, University of California San Diego, La Jolla, CA, USA.
[3]Department of Pharmacology, University of California San Diego, La Jolla, CA USA.
[4]Department of Chemistry and Biochemistry, University of California San Diego, La Jolla, CA, USA.
[5]Department of Medicine, University of California San Diego, CA, USA.
[6]UC San Diego HUMANOID™ Center of Research Excellence, San Diego, CA, USA.

† Equal contribution

**Conflict of interest statement:** Authors have declared that no conflict of interest exists.

**KEY WORDS**: Girdin, *CCDC88A*, atherosclerosis, macrophage, cholesterol transport, heterotrimeric G proteins

***Correspondence to**:

**Pradipta Ghosh, M.D.;** Professor, Departments of Medicine, and Cell and Molecular Medicine, University of California San Diego; 9500 Gilman Drive (MC 0651), George E. Palade Building, Rm 232, 239; La Jolla, CA 92093. Phone: 858-822-7633: Fax: 858-822-7636:

¶ **Alternative Address**: Department of General, Visceral and Transplant Surgery; University Hospital Tübingen; Germany.[i]

#### CATALOG OF SUPPLEMENTAL INFORMATION

1. *Supplemental Figures and Legends (S1-S10)*
2. *Supplemental Tables (S1-S3)*

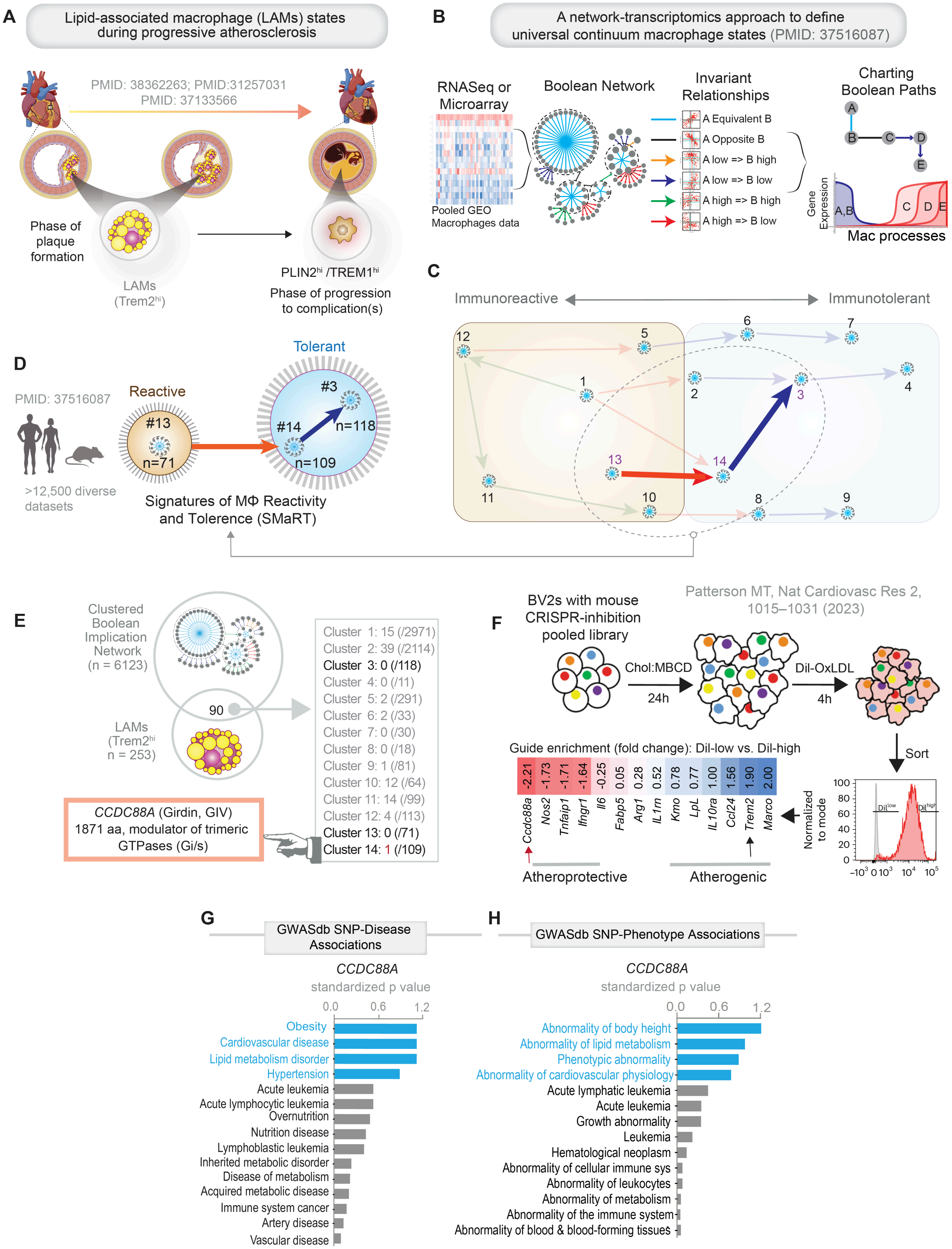


1 **Figure S1: Identification of *CCDC88A* as a prioritized target in lipid-associated macrophages (LAMs)**

**Related to Figure 1.**

**A**. Schematic summarizing scRNA-seq derived insights into lipid-associated macrophage (LAM) subpopulations that are enriched in atherosclerosis during its progression to complications (e.g., instability or rupture).

**B-C**. Key steps of the previously published workflow[1] used to develop the computational model of macrophage continuum states—Signatures of Macrophage Reactivity and Tolerance (SMaRT)—are shown (B). SMaRT identifies invariant gene clusters representing reactive (R) and tolerant (T1, T2) states across over ~12,500 diverse transcriptomic datasets (D and C). The schematic below illustrates their opposing roles: reactive macrophages, working antagonistically to fine-tune macrophage responses to the tissue environment (D and C).

**E**. Venn diagram showing overlap of the SMaRT signature (13-14-3) and a previously published[1] set of genes that are highly expressed and a distinguishing feature of TREM2hi LAMs. *CCDC88A* is identified as the only common gene. *See* **Data S1** for a complete set of SMaRT and LAM-defining genes.

**F**. Schematic showing the workflow for a previously published unbiased functional-genomics screen designed to identify regulators of macrophage lipid handling. pooled CRISPRi perturbations were introduced into BV2 macrophage-like cells, followed by DiI-LDL uptake assays to quantify lipid loading. Guide RNA enrichment analysis in DiI-low versus DiI-high populations enabled identification of genes with atheroprotective (red) or atherogenic (blue) functions[2].

**G-H**. GWASdb associations for *CCDC88A* with human diseases (G) and phenotypes (H) are shown. Blue bars highlight traits linked to cardiovascular and immunometabolic syndromes.

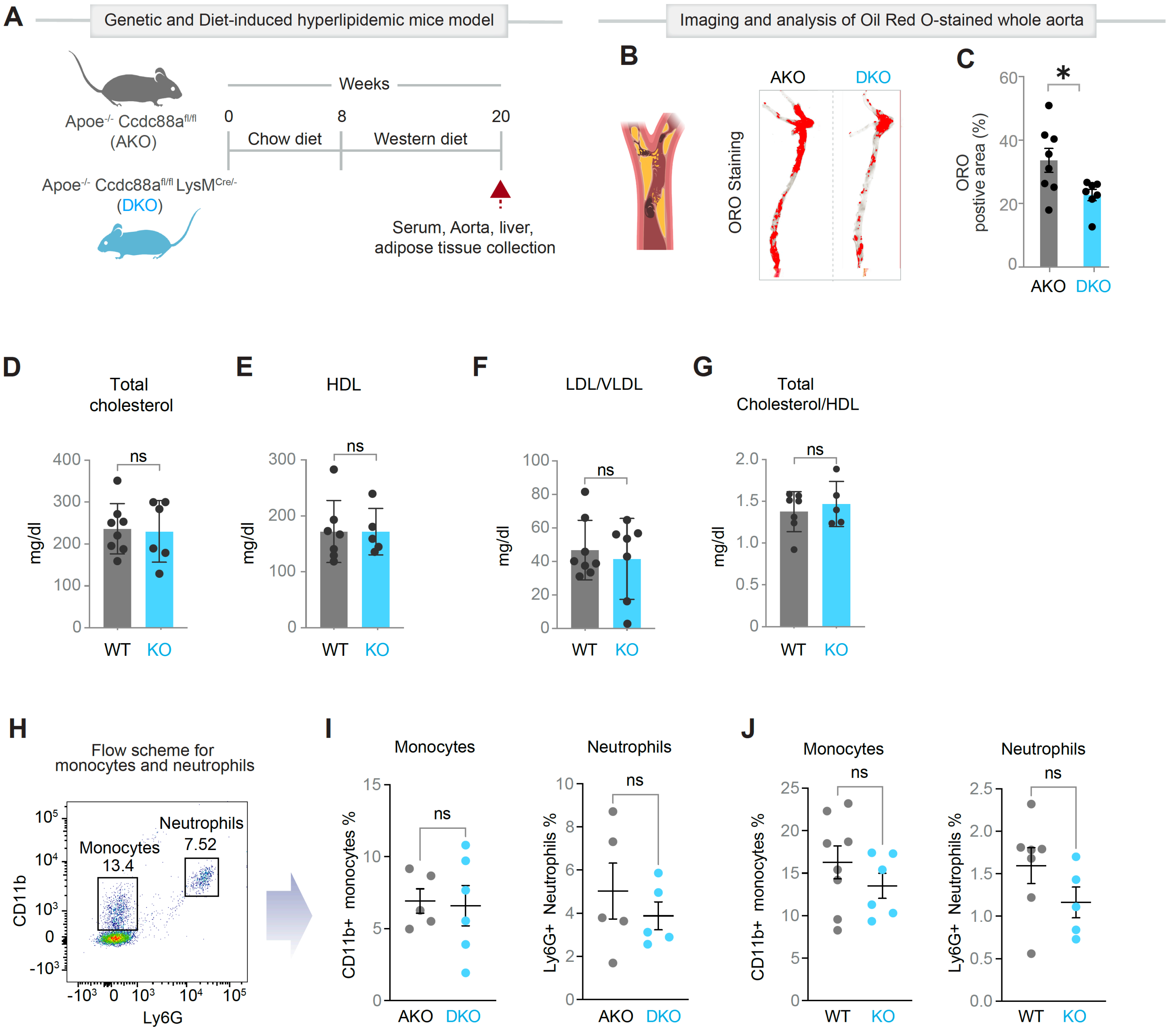


**Figure S2: Myeloid GIV loss reduces plaque burden without altering blood cholesterol and circulating myeloid populations**

**Related to Figure 2.**

**A.** Study design for the genetic and HFD-induced hyperlipidemic mice: 8-week-old mice were fed a HFD for 12 weeks prior to tissue collection (aorta, serum, and heart; n=7–8).

**B-C.** Representative *en face* aortic preparations stained with Oil Red O (B) and quantification of plaque lesion area (C, n=6-8). Scale bar, 2.5 mm.

**D-G.** Bar graphs showing serum lipid profiling: total cholesterol (D), HDL (E), LDL/VLDL (F), total-to-HDL cholesterol ratio (G). No significant differences observed.

**H-J.** Flow cytometry analysis of circulating leukocytes: gating strategy used to identify monocytes and neutrophils (H), and corresponding relative abundance in AKO vs DKO (I) and WT vs KO (J) shown in dot plots (I, K).

*Statistics*: All results are presented as mean ± SEM. Significance was assessed using an unpaired t-test. *p*-values < 0.05 were considered significant. $p < 0.05$ (*). ns = not significant.

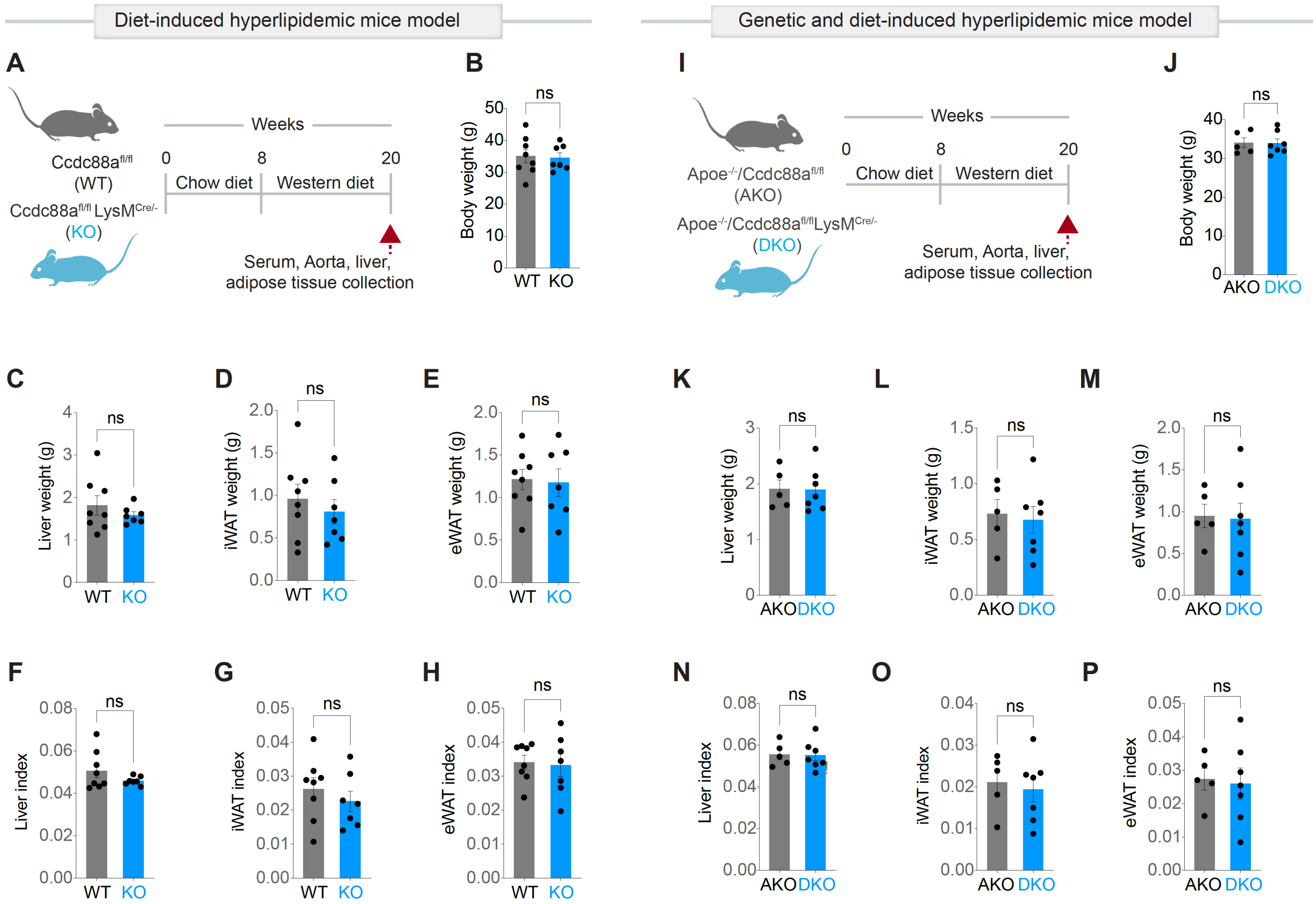


**Figure S3: GIV loss does not alter body, liver or adipose tissue mass in hyperlipidemic mouse models**

**Related to Figure 2.**

**A-H.** Study design (A) for the diet-induced hyperlipidemic (HFD; high-fat diet) mice model: 8-week-old WT and KO mice were fed a high-fat diet for 12 weeks prior to tissue collection (aorta, serum, and heart; n=7–8). Bar plots show body weight (B), liver weight (C), inguinal (i)WAT weight (D), epididymal (e)WAT weight (E), and corresponding organ-to-body-weight ratios (i.e., organ indices; F-H).

**I-P.** Study design (I) of genetic + HFD-induced hyperlipidemic mice model: 8 weeks old ApoE-KO (AKO) and double-KO (DKO; ApoE-KO + GIV-KO) mice were fed a HFD diet for 12 weeks prior to tissue collection (aorta, serum, and heart; n=7–8). Bar plots show body weight (J), liver weight (K), iWAT weight (L), eWAT weight (M), and corresponding organ indices (N-P).

*Statistics*: All results are presented as mean ± SEM. Significance was assessed using Welch's t-test. p-values < 0.05 were considered significant. ns = not significant.

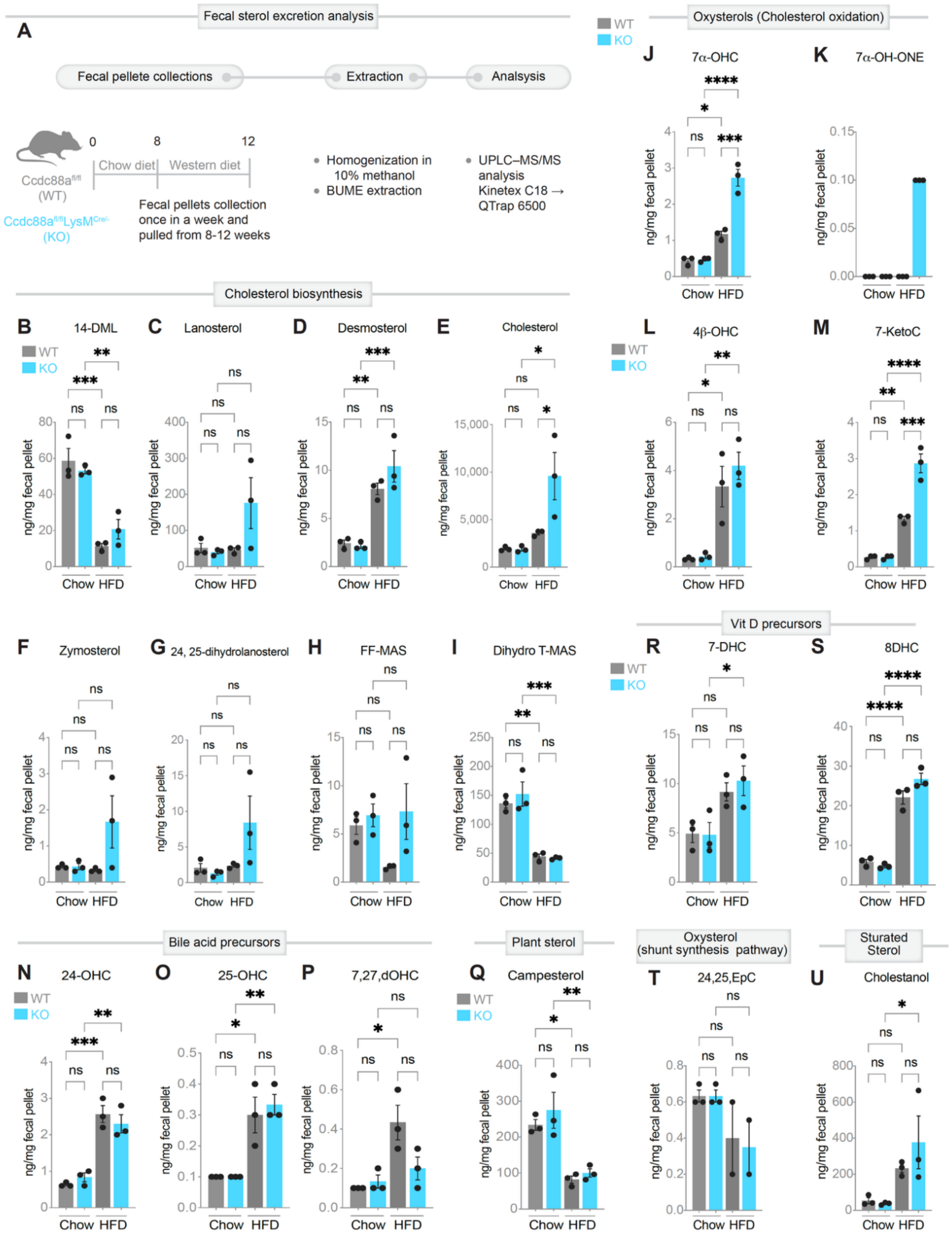


**Figure S4: Fecal sterol profiling during diet-induced hyperlipidemia**

**Related to Figure 2.**

**A.** Study design for the genetic and HFD-induced hyperlipidemic mice: 8-week-old mice were fed either chow diet or HFD. Fecal pellets were collected once every week and pooled from during weeks 4 - 8 of HFD exposure (n = 3). See also **Figure 2, Figure S2** and **Figure S3** for metabolic, hematologic and anatomical assessments, and **Figure S2A-C** for orthogonal validation in genetic (*ApoE*$^{-/-}$) hyperlipidemia model.
**B-U.** Quantification of fecal sterol species. Bar plots display concentrations of individual sterols associated with cholesterol absorption, metabolism, and microbial conversion (panel identifiers and sequence correspond to measurements referenced in **Figure 2K**).

*Statistics*: All results are presented as mean ± SEM. Significance was assessed using one-way ANOVA. *p*-values < 0.05 were considered significant. $p < 0.05$ (*), $p < 0.01$ (**), $p < 0.001$ (***) and $p < 0.0001$ (****).

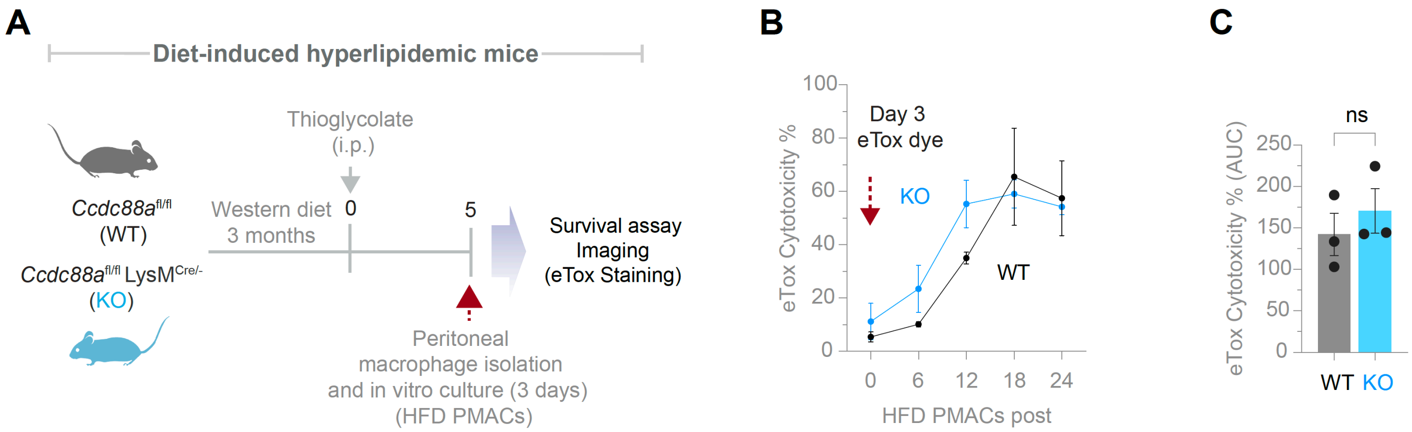


**Figure S5: Survival of macrophages from diet-induced hyperlipidemia**

**Related to Figure 3.**

**A.** Study design for diet-induced hyperlipidemia. Eight-week-old mice were fed a high-fat diet (HFD) for 12 weeks prior to macrophage isolation. Thioglycolate-elicited peritoneal macrophages (HFD PMACs) were harvested on day 5 following intraperitoneal injection of 2 ml of 3% thioglycolate, cultured in vitro for 3 days, and subsequently stained with eTOX cytotoxicity dye (n = 3). Live-cell imaging was performed using a BioTek Cytation system at 37°C and 5% $CO_2$ for 24 h.

**B-C.** Quantification of macrophage cytotoxicity. Line plots show percentage cytotoxicity measured at 6 h intervals over a 24 h period (B). Bar graphs display the corresponding area under the curve (AUC) for cytotoxicity kinetics shown in panel B (C).

*Statistics*: All results are presented as mean ± SEM. Statistical significance was determined using unpaired t-test or multiple unpaired t-tests, as appropriate. *p*-values < 0.05 were considered significant.

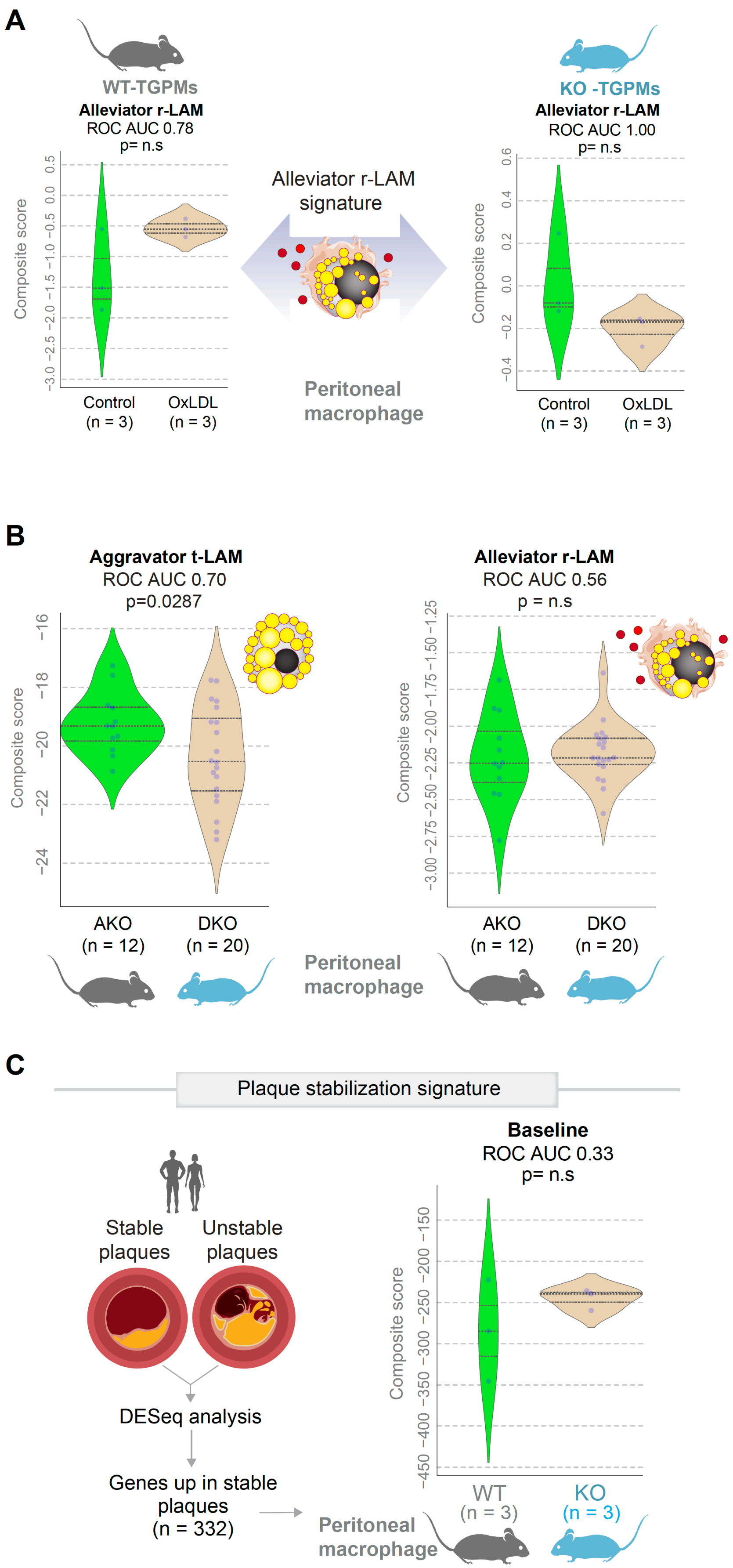


**Figure S6: Validation of lipid-associated macrophage (LAM) and plaque stabilization gene signatures**

**Related to Figure 4.**

**A**. Violin plots show composite scores for the alleviator reactive LAM (rLAM) signature (see **Figure 1C**) in WT and KO thioglycolate-elicited peritoneal macrophages (TGPMs), ± oxLDL stimulation. ROC AUC values indicate classification accuracy of the signature under each condition.

**B**. Composite scores for both aggravator tolerant LAM (tLAM) and alleviator reactive LAM (rLAM) signatures in orthogonal models of genetics + HFD-induced hyperlipidemia, AKO and DKO TGPMs, ± oxLDL exposure. ROC AUC values assess the discriminatory performance of signatures between genotypes and treatments.

**C.** A plaque-stabilization gene set (n = 332 genes; enlisted in **Data S5**), defined by DESeq2 analysis as upregulated in human stable plaques, was applied to TGPM transcriptomes. Violin plots display composite stabilization scores in WT versus KO conditions.

*Statistics*: All results are presented as mean ± SEM. Significance was assessed using Welch's t-test. p-values < 0.05 were considered significant. ns = not significant.

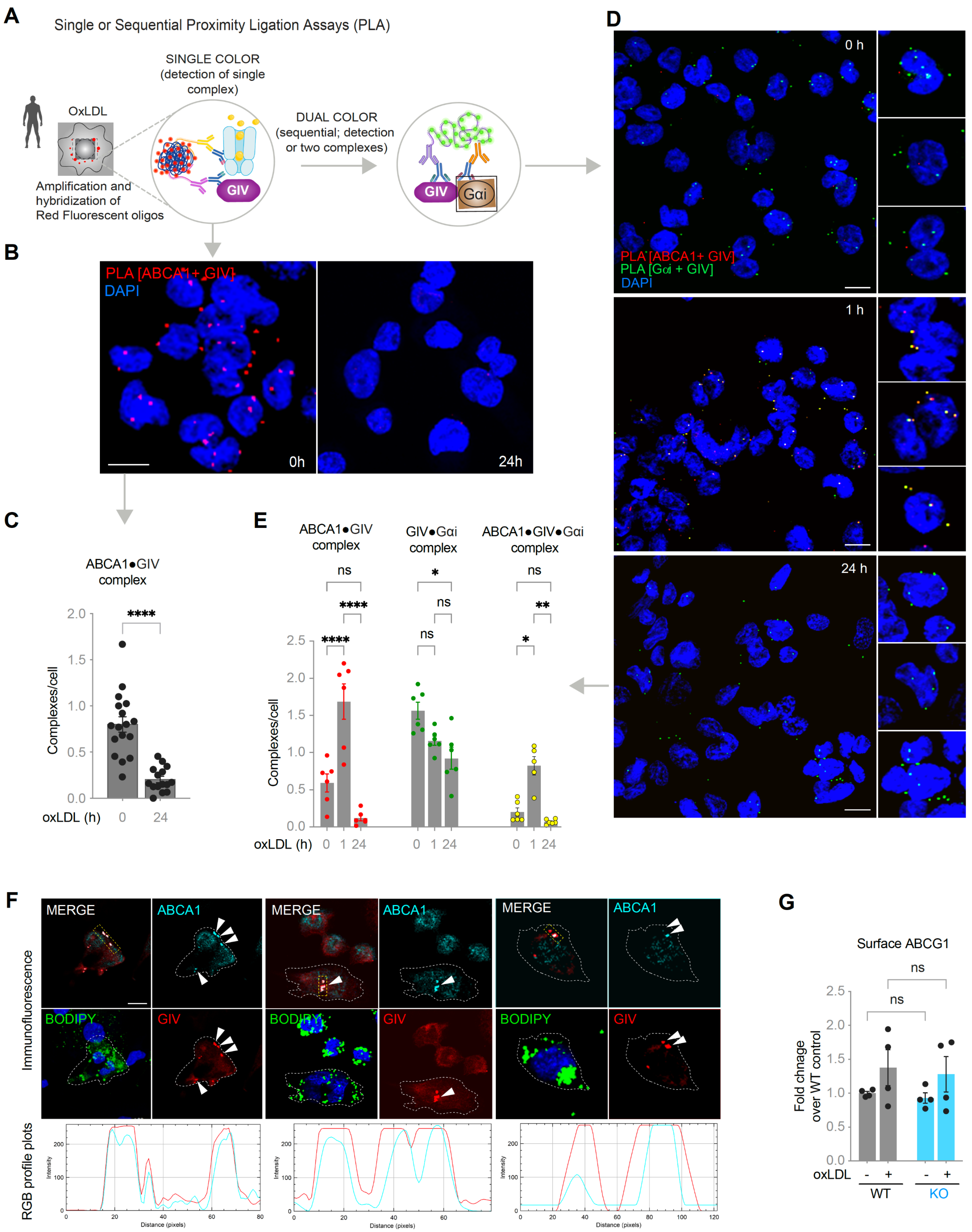


2 **Figure S7: GIV scaffolds ABCA1 to Gαi, sequestering the transporter in endomembranes**

3 **Related to Figure 5.**

**A**. Workflow for *in situ* proximity ligation assays (PLA), implemented either as single- or sequential dual-color, to visualize and quantify ABCA1-bound complexes (B-L) in WT and GIV-KO THP1-derived macrophages.
**B-C**. Dynamics of ABCA1●GIV complexes upon oxLDL (100 µg/ml) treatment detected by single-color PLA. Representative images (B) and quantification (C). Scale bar, 10 µm.
**D-E**. ABCA1●GIV●Gαi ternary complexes detected by dual-color PLA. Representative images (D) and quantification (E). Scale bar, 10 µm.
**F**. Representative montage of immunofluorescence micrographs of WT THP1-derived macrophages showing subcellular co-localization of ABCA1 (cyan) and GIV (red) in lipid droplet–laden LAMs, as determined by BODIPY (green). Of note, colocalization of ABCA1 and GIV did not occur on lipid droplets. Line-scan intensity plots (bottom) correspond to boxed regions (top).
**G**. Surface ABCG1 measured by flow cytometry and quantified using NovoExpress software.

*Statistics*: All results are presented as mean ± SEM. Significance was assessed using one-way ANOVA. *p*-values $< 0.05$ were considered significant. $p < 0.05$ (*), $p < 0.01$ (**), $p < 0.001$ (***) and $p < 0.0001$ (****).

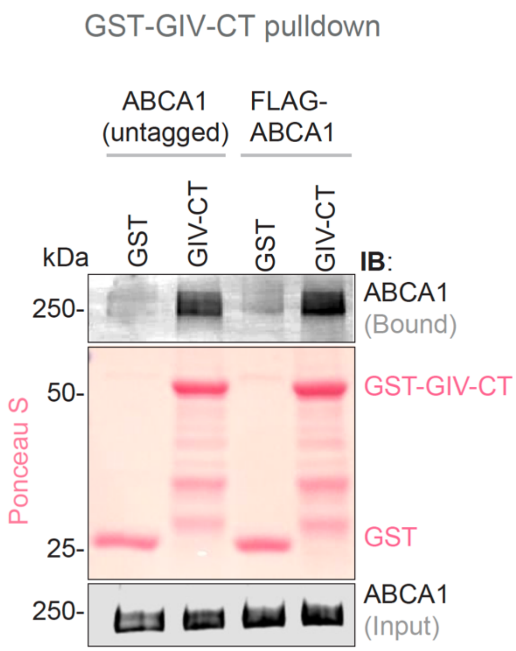


**Figure S8: GIV's C-terminus binds to full-length ABCA1**

**Related to Figure 6.**

GST pulldown assays using bead-immobilized GST–GIV-CT and lysates of HEK293T cells exogenously expressing full-length untagged or FLAG-ABCA1, probed for ABCA1 binding. Equal loading of GST proteins is confirmed by Ponceau S staining.

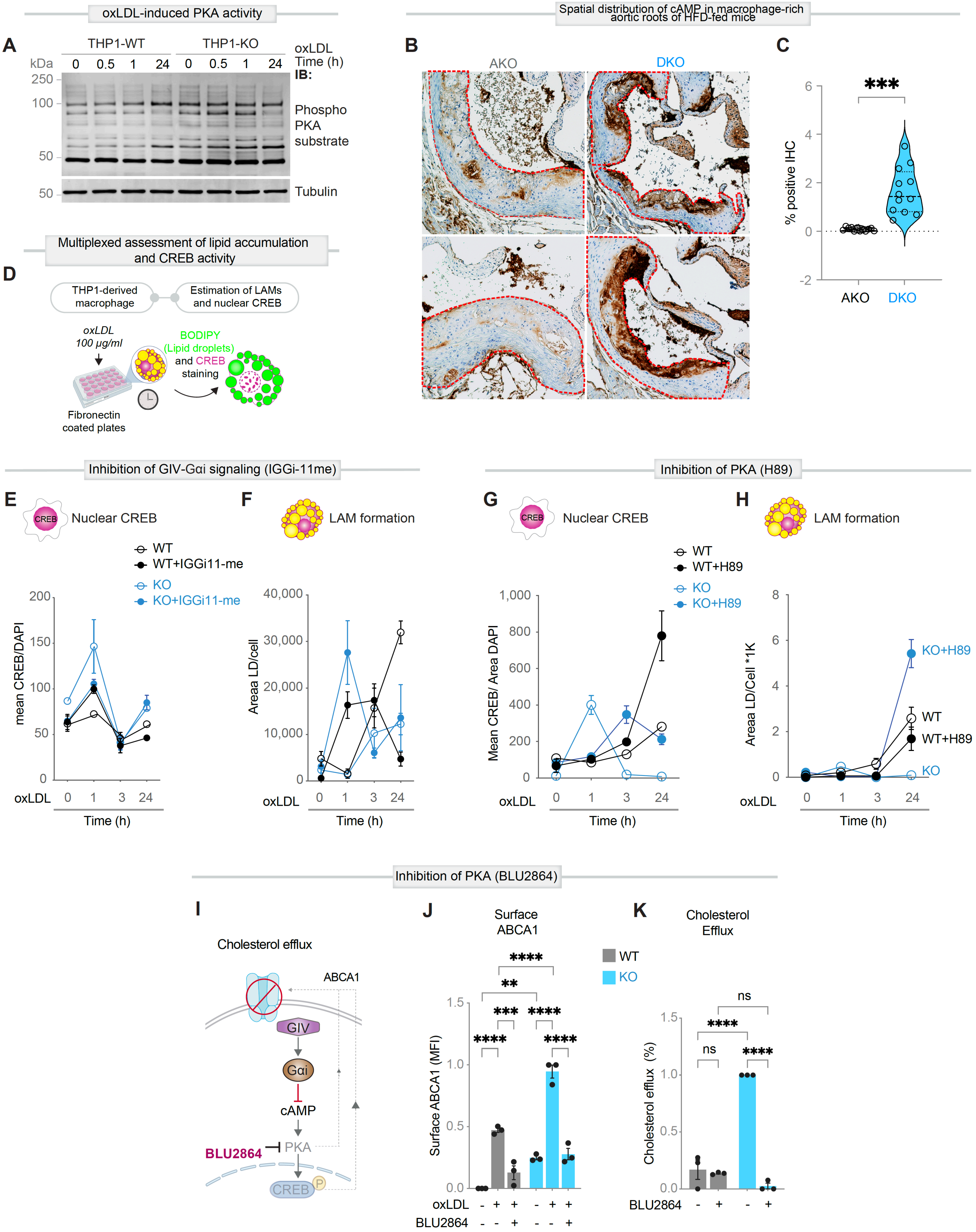


**Figure S9: Pharmacogenomic evaluation of how the PKA-CREB pathway impacts LAM formation**

**Related to Figure 7.**

**A.** Immunoblot of WT and GIV-KO THP1-derived macrophages stimulated with oxLDL (100 µg/ml, 0-8 h), assessing PKA activity at the indicated time points, as determined using a phospho-substrate antibody.

**B-C.** Macrophage-specific deletion of GIV increases cAMP in macrophage-rich aortic root regions. Representative immunohistochemical images showing the spatial distribution of cAMP in macrophage-rich aortic root sections from HFD-fed AKO and DKO mice. Red dashed outlines demarcate the boundaries of atherosclerotic lesions within the aortic root (B). Quantification of cAMP-positive area within aortic root lesions. Each dot represents an individual aortic lesion (C).

**D-H**. Workflow schematic (D) for inducing LAM formation in oxLDL-injured THP1-derived macrophages while simultaneously assessing cells for CREB signals, as determined by its nuclear localization. plots show the impact of GIV●Gαi inhibition (10 µM, IGGi-11me) on nuclear CREB activation (E) and LAM differentiation (F). Complementary studies using a PKA inhibitor (30 µM, H89) evaluate effects on nuclear CREB (G) and LAM formation (H).

**I-K**. PKA inhibition enforces foam cell fate: Schematic of BLU2864 intervention (10 µM) and its effect on ABCA1 cell-surface localization (J) and cholesterol efflux function (K) in WT and GIV-KO THP1-derived macrophages.

*Statistics*: All results are presented as mean ± SEM. Significance was assessed using an unpaired t-test or one-way ANOVA. *p*-values < 0.05 were considered significant. $p < 0.05$ (*), $p < 0.01$ (**), $p < 0.001$ (***) and $p < 0.0001$ (****).

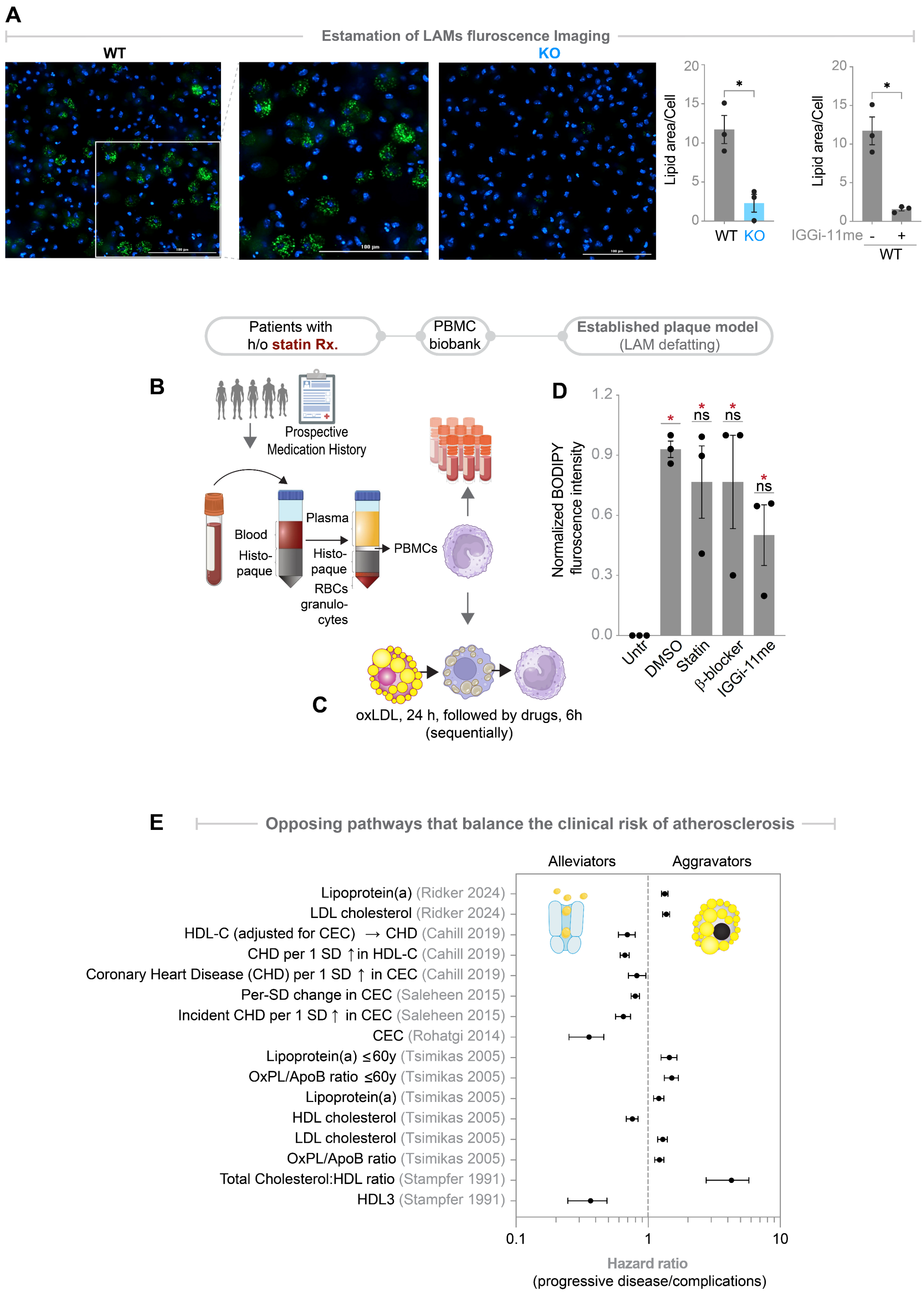


**Figure S10: Pharmacologic disruption of the GIV–Gαi checkpoint reverses macrophage lipid loading and restores cholesterol efflux programs, counterbalancing pathways in atherosclerosis risk**

**Related to Figure 8.**

**A**. *Ex vivo* analysis of lipid-loading in LAMs (See **Figure 8A** for study design). Eight-week-old mice were maintained on high-fat diet (HFD) for 3 months, followed by thioglycolate elicitation and intraperitoneal administration of IGGi-11me (1mg/kg), followed by isolation of peritoneal macrophages (HFD-PMACs) for lipid quantification by fluorescence imaging (n=3). Representative images (left) and quantification of lipid droplet area per cell (right) demonstrate marked lipid accumulation in WT, but not GIV-KO, macrophages. Scale bar, 100 µm.
**B-D**. Schematic of workflow for therapeutic testing in PBMC-derived macrophages on patients with systemic lipid overload under treatment with statins, as mimicked by challenging them with 100 µg/ml oxLDL (A). Compounds were added after LAM maturation to test their "defatting" (reversal of injury) capacity (B). Bar graphs show the effects of statin (10 µM), β-blocker (10 µM), and IGGi-11me (10 µM) on defatting LAMs (C).
**E**. Forest plot summarizing hazard ratios (median with 95% confidence intervals) for lipid-related factors associated with the clinical risk of atherosclerosis. LipoA, LDL cholesterol and other surrogate measures of oxidized cholesterol post as aggravator of disease (right side), whereas cholesterol efflux capacity (CEC) or HDL, which is a byproduct of CEC, are alleviators of risk.

*Statistics*: Data shown as mean ± SEM. Significance was assessed by one-way ANOVA. Significance: $p < 0.05$. ns = not significant.

### SUPPLEMENTAL TABLES

**Table S1: List of genes in the LAM gene signature, related to Figure 1B**

| Alleviator reactive rLAMs gene signature (4 genes) | Aggravator tolerant tLAMs gene signature (20 genes) |
|---|---|
| LIMK2 | **CCDC88A** |
| SFT2D2 | HNMT |
| STAT3 | KMO |
| KIAA0040 | MGAT4A |
| | CSE1L |
| | CLEC4A |
| | RGS10 |
| | CREBL2 |
| | NOL8 |
| | F13A1 |
| | TCN2 |
| | ZNF589 |
| | MS4A6A |
| | HEXA |
| | CCDC93 |
| | CPVL |
| | DNASE1L3 |
| | UBL3 |
| | ARL4C |
| | UAP1L1 |

**Table S2: The odds ratio of atherosclerosis based on clinical and molecular gene signatures, related to Figure 8I-K**

| Parameter | Bad + | Bad - | Good + | Good - | 95CL low | Odds Ratio | 95CL High | *p*-value | 95% CI (low–high) |
|---|---|---|---|---|---|---|---|---|---|
| **Sex (Male)** | 10 | 5 | 8 | 7 | 0.4 | 1.75 | 7.66 | 0.71 | 0.40 – 7.66 |
| **Smoker** | 7 | 8 | 7 | 8 | 0.24 | 1 | 4.2 | 1 | 0.24 – 4.20 |
| **Hypertension** | 14 | 1 | 10 | 5 | 0.71 | 7 | 69.49 | 0.17 | 0.71 – 69.49 |
| **Hyperlipidemia** | 6 | 9 | 7 | 8 | 0.18 | 0.76 | 3.24 | 1 | 0.18 – 3.24 |
| **β-blockers** | 5 | 10 | 6 | 9 | 0.17 | 0.75 | 3.33 | 1 | 0.17 – 3.33 |
| **Aspirin/Clopidogrel** | 12 | 3 | 14 | 1 | 0.03 | 0.29 | 3.12 | 0.6 | 0.03 – 3.12 |
| **Statins** | 9 | 6 | 9 | 6 | 0.23 | 1 | 4.31 | 1 | 0.23 – 4.31 |
| **Thrombosis prophylaxis** | 5 | 10 | 8 | 7 | 0.1 | 0.44 | 1.92 | 0.46 | 0.10 – 1.92 |
| **ACE-inhibitor** | 7 | 8 | 6 | 9 | 0.31 | 1.31 | 5.58 | 1 | 0.31 – 5.58 |
| **Aggravator tLAM** | 12 | 4 | 2 | 11 | 2.51 | 16.5 | 108.6 | 0.0025 | 2.51-108.60 |
| **RCT** | 2 | 14 | 12 | 1 | 0.001 | 0.012 | 0.148 | 0.000021 | 0.001-0.148 |

**Table S3: The odds ratios for atherosclerosis based on risk factors, related to Figure 8 and S10.**

| Study | HR | CI Lower | CI Upper | Authors | Journal/Year |
|---|---|---|---|---|---|
| **HDL3** | 0.3 | 0.2 | 0.6 | Stampfer M et al., [3] | N Engl J Med 1991 |
| **Total Cholesterol:HDL ratio** | 3.73 | 1.95 | 7.12 | Stampfer M et al., [3] | N Engl J Med 1991 |
| **OxPL/apoB ratio** | 1.21 | 1.05 | 1.39 | Tsimikas S et al.,[4] | N Engl J Med 2005 |
| **LDL cholesterol** | 1.28 | 1.11 | 1.48 | Tsimikas S et al., [4] | N Engl J Med 2005 |
| **HDL cholesterol** | 0.75 | 0.63 | 0.9 | Tsimikas S et al., [4] | N Engl J Med 2005 |
| **Lipoprotein(a)** | 1.2 | 1.02 | 1.4 | Tsimikas S et al., [4] | N Engl J Med 2005 |
| **OxPL/apoB ratio ≤60y** | 1.49 | 1.2 | 1.84 | Tsimikas S et al., [4] | N Engl J Med 2005 |
| **Lipoprotein(a) ≤60y** | 1.42 | 1.12 | 1.81 | Tsimikas S et al., [4] | N Engl J Med 2005 |
| **Cholesterol efflux capacity** | 0.33 | 0.19 | 0.55 | Rohatgi A et al.,[5] | N Engl J Med 2014 |
| **Incident CHD per 1 SD ↑ in CEC** | 0.64 | 0.51 | 0.8 | Saleheen et al.,[6] | - |
| **Per-SD change in CEC** | 0.8 | 0.7 | 0.9 | Saleheen et al.,[6] | - |
| **Coronary Heart Disease (CHD) per 1 SD ↑ in CEC** | 0.82 | 0.71 | 0.96 | Cahill LE et al.,[7] | J Lipid Res, 2019 |
| **CHD per 1 SD ↑ in HDL-C** | 0.66 | 0.58 | 0.76 | Cahill LE et al., [7] | - |
| **HDL-C (adjusted for CEC) → CHD** | 0.68 | 0.53 | 0.88 | Cahill LE et al., [7] | - |
| **LDL cholesterol** | 1.36 | 1.23 | 1.52 | Ridker P.M et al.,[8] | N Engl J Med 2024 |
| **Lipoprotein(a)** | 1.33 | 1.21 | 1.47 | Ridker P.M et al.,[8] | N Engl J Med 2024 |
